# Distinct Colon Mucosa Microbiomes associated with Tubular Adenomas and Serrated Polyps

**DOI:** 10.1101/2021.07.20.453135

**Authors:** Julio Avelar-Barragan, Lauren DeDecker, Zachary Lu, Bretton Coppedge, William E. Karnes, Katrine L. Whiteson

## Abstract

**Background:** Colorectal cancer is the second most deadly and third most common cancer in the world. Its development is heterogenous, with multiple mechanisms of carcinogenesis. Two distinct mechanisms include the adenoma-carcinoma sequence and the serrated pathway. The gut microbiome has been identified as a key player in the adenoma-carcinoma sequence, but its role in serrated carcinogenesis is less clear. In this study, we characterized the gut microbiome of 140 polyp-free and polyp-bearing individuals using colon mucosa and fecal samples to determine if microbiome composition was associated with each of the two key pathways.

**Results:** We discovered significant differences between the microbiomes of colon mucosa and fecal samples, with sample type explaining 14% of the variation observed in the microbiome. Multiple mucosal samples were collected from each individual to investigate whether the gut microbiome differed between polyp and healthy intestinal tissue, but no differences were found. Colon mucosa sampling revealed that the microbiomes of individuals with tubular adenomas and serrated polyps were significantly different from each other and polyp-free individuals, explaining 2-10% of the variance in the microbiome. Further analysis revealed differential abundances of 6 microbes and 1,143 microbial genes across tubular adenoma, serrated polyp, and polyp-free cases.

**Conclusion:** By directly sampling the colon mucosa and distinguishing between the different developmental pathways of colorectal cancer, this study helps characterize potential mechanistic targets for serrated carcinogenesis. This research also provides insight into multiple microbiome sampling strategies by assessing each method’s practicality and effect on microbial community composition.

## Introduction

Colorectal cancer (CRC) is the second most deadly and third most common cancer globally, accounting for over 900,000 deaths in 2020.^1^ The etiologies of CRC are multifactorial, with only 5-10% of cases being attributable to hereditary germline mutations.^2^ Significant risk factors include diets high in red meat and low in fiber, obesity, physical inactivity, drug and alcohol usage, and chronic bowel inflammation.^3–6^ Each of these factors is associated with compositional and functional changes in the collective community of bacteria, fungi, archaea, and viruses that inhabit the colon.^7–10^ Commonly referred to as the gut microbiome, this community of microorganisms has been identified as a potential regulator of CRC initiation and progression.

Colorectal polyp formation precedes cancer development and is influenced by various environmental factors and host genetics. Polyps most commonly progress into malignancy through the adenoma-carcinoma sequence.^11^ This pathway is characterized by chromosomal instability and mutations in the adenomatous polyposis coli (APC) gene, KRAS oncogene, and TP53 tumor suppressor gene.^12^ Alternatively, 15 to 30% of CRCs develop through the serrated pathway.^13^ This pathway is characterized by the epigenetic hypermethylation of gene promoters to produce a CpG island methylator phenotype.^13^ In addition to the epigenetic inactivation of tumor suppressor genes, BRAF or KRAS mutations are also common.^13^ The serrated pathway often results in the production of hyperplastic polyps (HPPs), traditional serrated adenomas (TSAs), and sessile serrated polyps (SSPs).^14^ Premalignant polyps from both pathways can be screened for and removed during colonoscopy to prevent CRC formation, but incomplete polyp resection or escaped detection can result in the development of interval cancers. Compared to other colorectal polyps, SSPs are disproportionately responsible for interval cancers, as their flat morphology makes them difficult to detect.^15^ Additional detection methods, such as SSP-specific biomarkers, would assist with CRC prevention.

One potential avenue for polyp-specific biomarker discovery is the gut microbiota. SSPs often overexpress mucin forming genes, like MUC6, MUC5aC, MUC17, and MUC2, producing a mucus cap, which may harbor unique, mucin-degrading microbes.^16^ Finding microbiome alterations in patients consistent with the presence of SSPs would enable gastroenterologists to personalize their technique and screening frequency for these higher risk patients. Additionally, elucidating the microbiome alterations specific to the adenoma-carcinoma sequence or the serrated pathway would help better understand the mechanisms of how particular microbes, their metabolites, and dysbiosis may contribute to colorectal carcinogenesis.

Studies comparing the microbiomes of these two pathways with healthy controls have yet to discover differences between healthy individuals and those with serrated polyps.^17–19^ One reason for this may be the dominant use of stool for characterizing the microbiome, which does not accurately represent microbes adherent to the intestinal epithelium.^20, 21^ In this regard, we hypothesized that colon mucosa samples would more accurately reflect the composition of microbes intimately associated with colorectal polyps. To investigate this, and the role of the microbiome in the adenoma-carcinoma and serrated pathways, we used multiple sampling techniques to obtain microbiome samples from colorectal polyps. Samples were collected during and after colonoscopy from healthy individuals or those with tubular adenomas (TAs), HPPs, or SSPs. When possible, samples from the same individual were collected from polyps and the healthy colon epithelium opposite from these polyps. Stool samples were also collected 4-6 weeks after colonoscopy. We used a combination of amplicon (16S and ITS) and shotgun sequencing to characterize the microbial communities of samples. The purpose of our work was to 1) develop and compare microbiome sampling methods during colonoscopy; 2) determine whether microbiomes differ between polyp and healthy tissue samples within the same individual; and 3) identify microbiome members or genes specific to CRC precursors in the adenoma-carcinoma sequence versus the serrated pathway. Our key hypothesis was that there would be distinct differences between the microbiomes of individuals with tubular adenomas versus serrated polyps.

## Methods and Materials

### Subject Recruitment and Criteria

Individuals who presented for colonoscopy with indications of screening for, or a prior history of, colorectal polyps were asked to participate in the study. Written and informed consent was obtained from each subject and was required for participation. Subjects who were pregnant, had taken antibiotics within 6 weeks of colonoscopy, or with known inflammatory bowel diseases, were excluded. In total, 140 individuals were recruited for this study. Of the 140 individuals, 50 were found to be polyp-free, 45 had one or more TAs, 33 had polyps originating from the serrated pathway (HPPs and SSPs), and 12 had unknown or other pathologies (Figure 1).

**Figure 1:**
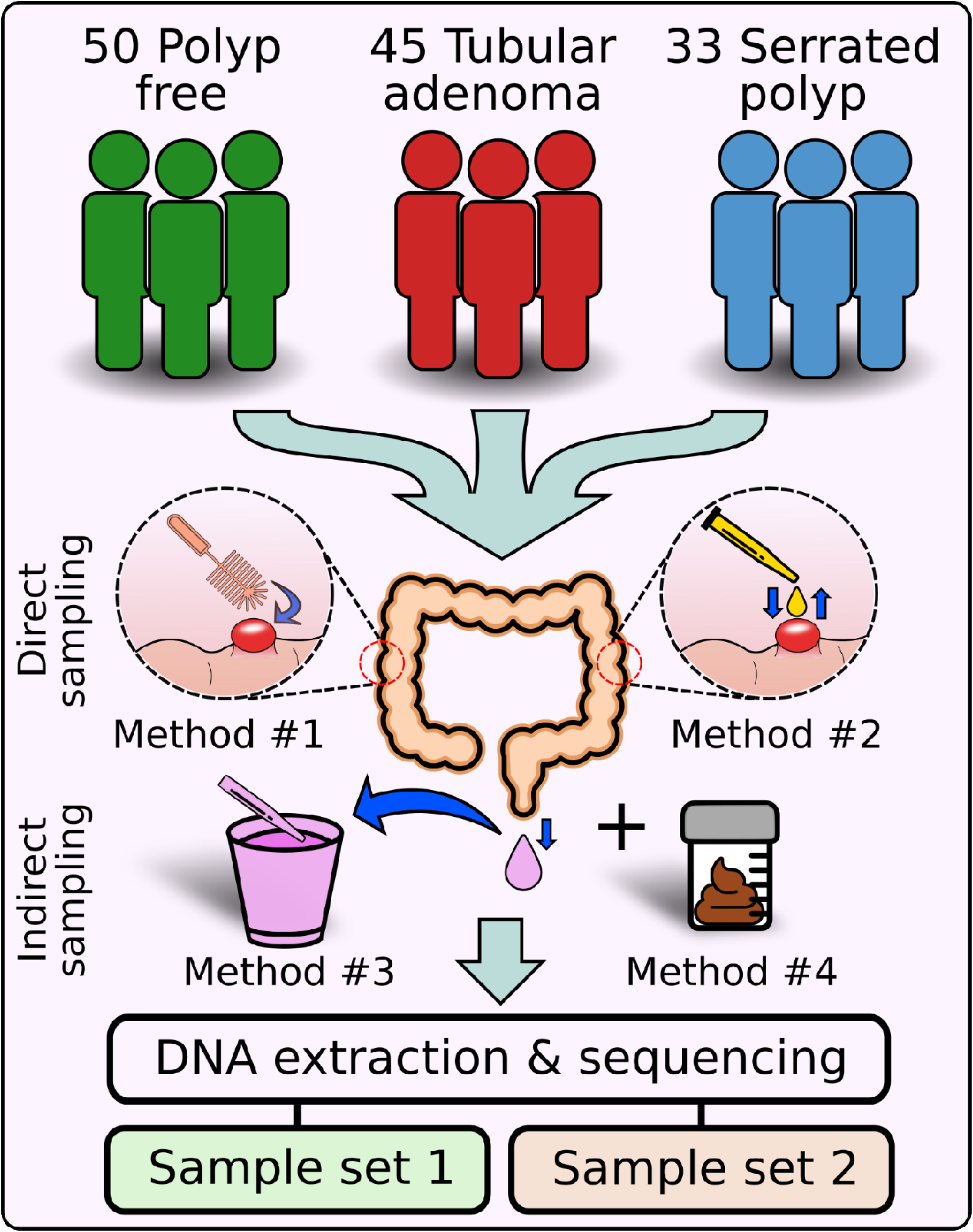
Study design. A total of 140 individuals were recruited for this study, including 50 polyp-free individuals, 45 with tubular adenomas, and 33 with serrated polyps (HPP, TSA, or SSP). For the remaining 12 individuals, 11 had unknown pathology and one had an adenocarcinoma. Multiple samples were taken from each subject during colonoscopy. This included mucosal brushes (Method #1, orange), mucosal aspirates (Method #2, yellow), and lavage aliquots (Method #3, purple). Fecal samples (Method #4, brown) were collected from participants four to six weeks post-colonoscopy. DNA extraction and sequencing produced two sample sets. The first sample set was produced by sequencing mucosal brushes, mucosal aspirates, and lavage aspirates using 16S and ITS sequencing. The second sample set was produced by sequencing mucosal aspirates, lavage aspirates, and fecal samples using whole-genome shotgun sequencing.

### Colonoscopy Preparation, Procedure, and Sample Collection

Before colonoscopy, subjects were asked to adhere to a clear liquid diet for 24 hours. Bowel cleansing was done using Miralax, or polyethylene glycol with electrolytes administered as a split dose, 12 and 5 hours before the procedure. Sample collection focused on two direct and two indirect microbiome sampling methods (Figure 1). The first direct sampling method involved brushing the mucosa of colon tissue during colonoscopy (Method #1 in Figure 1). Since mucosal brushes can potentially damage or agitate the intestine, we also employed a novel method of direct microbiome sampling in which colonoscopy washing fluid was sprayed directly on to the target mucosa and immediately re-suctioned into a storage vial (Method #2 in Figure 1). Participants with suspected polyps had additional mucosal washing aspirates taken near the polyp, as well as brushings of the polyp and opposite wall of the polyp to study the microbial microenvironment. When more than one polyp was found, the largest polyp was targeted for mucosal brushing and aspirate sampling. The first indirect sampling method involved collecting an aliquot of the post-colonoscopy lavage fluid (Method #3 in Figure 1). This lavage fluid was produced from rinsing the wall of the colon throughout the procedure and was collected in a catch can outside the subject. All samples were collected in sterile cryogenic tubes and placed on ice until the colonoscopy procedure was finished. Afterwards, the samples were stored at -80°C. Additional information collected included indication for procedure, age, sex, ethnicity, BMI, family history, and findings, including the size, shape, location, and pathology of all polyps sampled.

### Patient-directed Collection of Fecal Samples

For the second indirect microbiome sampling method, subjects were encouraged to send follow-up fecal samples four to six weeks post-colonoscopy (Method #4 in Figure 1). Subjects were provided with a fecal collection kit, which contained collection equipment, prepaid shipping labels, and Zymo DNA/RNA shield preservation buffer (R1101). Subjects who complied were compensated $20 USD. Samples were returned via the United States Postal Service. After arrival, samples were stored at -80°C. Thirty-eight fecal samples were returned, bringing our total number of samples collected to 1,883. A summary of the sample types can be found in Supplemental Table 1.

### Polyp and Subject Type Classification

Polyp biopsies collected during colonoscopy were sent to a pathologist for classification. This information was then recorded for the corresponding mucosal brush aspirate samples. Pathology reports were also used to broadly categorize all samples collected from an individual by their polyp pathology. We referred to this as the ‘subject type’ and the three categories were polyp-free subjects, TA-bearing subjects, and serrated polyp-bearing (SP-bearing) subjects, which included both HPPs and SSPs. For example, if a sample was taken from healthy intestinal tissue of an individual who was found to have a TA, that sample and all others from the same individual would be included in the TA-bearing subject type. Three individuals had both a TA and an SSP and were classified as SP-bearing subjects.

### DNA Extraction

Two separate DNA extractions were performed in this study, yielding two different sample sets (Table 1). Sample set 1 DNA extractions included mucosal brushes, mucosal aspirates, and lavage aspirates only. Sample set 2 DNA extractions occurred later and included mucosal aspirates, lavage aspirates, and fecal samples. All samples were thawed on ice for DNA extraction. For mucosal aspirates and lavage aliquot samples, 250 uL of fluid were taken from each sample and then DNA was extracted using ZymoBiomics DNA Miniprep Kit (D4300) according to the manufacturer’s protocol. For mucosal brushes, 750 uL of ZymoBIOMICS Lysis Solution was mixed with the brushes in their original sterile cryogenic tube and vortexed for 5 minutes to suspend the contents of the brush into solution. The solution was then transferred and extracted according to the manufacturer’s protocol. Fecal samples stored in Zymo DNA/RNA shield were thawed, mixed by vortexing, and 750 uL of the fecal plus buffer mix was extracted according to the manufacturer’s protocol.

**Table 1:**
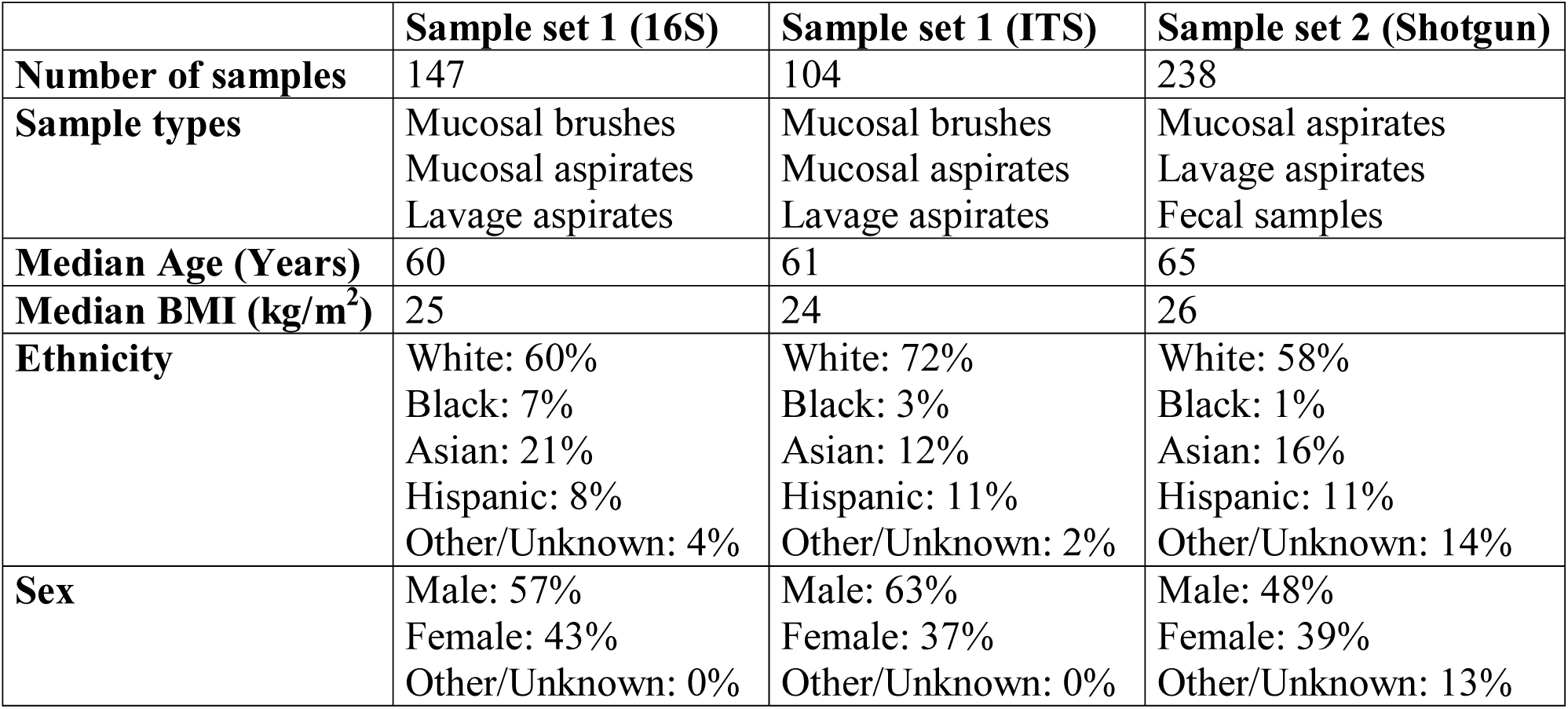
***Study cohort information.*** A table describing the sample sizes, sample types, median age, median BMI, ethnicity compositions, and sex ratios of each sample set. The first sample set was sequenced twice, once using 16S sequencing and once using ITS sequencing.

|  | **Sample set 1 (16S)** | **Sample set 1 (ITS)** | **Sample set 2 (Shotgun)** |
| --- | --- | --- | --- |
| **Number of samples** | 147 | 104 | 238 |
| **Sample types** | Mucosal brushes<br>Mucosal aspirates<br>Lavage aspirates | Mucosal brushes<br>Mucosal aspirates<br>Lavage aspirates | Mucosal aspirates<br>Lavage aspirates<br>Fecal samples |
| **Median Age (Years)** | 60 | 61 | 65 |
| **Median BMI (kg/m<sup>2</sup>)** | 25 | 24 | 26 |
| **Ethnicity** | White: 60%<br>Black: 7%<br>Asian: 21%<br>Hispanic: 8%<br>Other/Unknown: 4% | White: 72%<br>Black: 3%<br>Asian: 12%<br>Hispanic: 11%<br>Other/Unknown: 2% | White: 58%<br>Black: 1%<br>Asian: 16%<br>Hispanic: 11%<br>Other/Unknown: 14% |
| **Sex** | Male: 57%<br>Female: 43%<br>Other/Unknown: 0% | Male: 63%<br>Female: 37%<br>Other/Unknown: 0% | Male: 48%<br>Female: 39%<br>Other/Unknown: 13% |

### 16S Amplicon Library Preparation and Sequencing

Samples from the first set underwent 16S and ITS amplicon sequencing. We targeted the V4 region of the bacterial 16S rRNA gene using the 515F and 926R primers. For each sample, the V4 region was amplified using 25 uL polymerase chain reaction (PCR) volumes with the following reagents: 12.5 uL of 1x AccustartII PCR tough mix (QuantaBio 95142), 9.5 uL of PCR grade water, 1 uL of 10 mg/mL BSA, 0.5 ng of extracted genomic DNA, and 0.5 uL of 0.2 uM 515F and 926R primers each. The 515F primer contained the Illumina adapter sequence and barcode. Each sample was amplified using a thermocycler for 30 cycles (94°C for 3 min; 94°C for 45 sec, 55°C for 30 sec, 72°C for 20 sec; repeat steps 2-4 30 times; 72°C for 10 min). The resultant amplicons were quantified using the Qubit dsDNA HS Assay Kit (Life technologies Q32851) according to the manufacturer’s protocol and pooled at equimolar concentrations. The pooled amplicon library was cleaned and concentrated using Agencourt AMPure XP beads (Beckman-Coulter A63880) according to the manufacturer’s protocol. Equimolar PhiX was added at 10% final volume to the amplicon library and sequenced on the Illumina MiSeq platform, yielding 300bp paired-end sequences. An average of 41,578 +/- 35,920 (σ) reads per sample were obtained for 16S amplicons.

### ITS Amplicon Library Preparation and Sequencing

Fungi from the first sample set were characterized by targeting the ITS2 region of the 18S rRNA gene for amplification. We used the ITS9f and ITS4r primers, as described by Looby et al.^22^ PCR was performed in 25 uL volumes, consisting of: 12.5 uL of 1x AccustartII PCR tough mix, 9.5 uL of PCR grade water, 1 uL of 10 mg/mL BSA, 0.5 ng of extracted genomic DNA, and 0.5 uL of 0.3 uM ITS9f and barcoded ITS4r primers each. Amplification was performed with the following thermocycler settings: 94°C for 5 min, 35 cycles of 95°C for 45 sec, 50°C for 1 min, 72°C for 90 sec, and a final extension step of 72°C for 10 min. Afterwards, we quantified, pooled, and cleaned our ITS2 amplicons using the same methods as our 16S amplicons. Our ITS2 library was combined with our 16S library and sequenced simultaneously in the reverse complementary orientation. This yielded an average of 22,252 +/- 17,000 (σ) ITS reads per sample.

### Shotgun Library Preparation and Sequencing

The second sample set was sequenced using shotgun sequencing. Libraries were prepared using the Illumina DNA prep kit (20018705), using our low-volume protocol.^23^ Briefly, a maximum of 5 uL or 50 ng (whichever was reached first) of DNA from each sample was tagmented with 2 uL of tagmentation master mix for 15 min at 55°C. Afterwards, 1 uL of tagmentation stop buffer was added to each sample and incubated at 37°C for 15 min. The samples were washed with the provided buffer according to the manufacturer’s protocol, then PCR was performed with 12.5 uL reaction volumes with the following reagents: 6.25 uL of KAPA HiFi HotStart ReadyMix (Roche Life Science KK2602), 2.75 uL of PCR grade water, 1.25 uL of 1 uM i5 and i7 index adaptors each, and 0.5 uL of 10 uM forward and reverse KAPA HiFi polymerase primers each. PCR amplification was done with the settings: 72°C for 3 min, 98°C for 3 min, 12 cycles of 98°C for 45 sec, 62°C for 30 sec, 72°C for 2 min, and a final extension step of 72°C for 1 min. Samples were pooled and size selection was performed per the manufacturer’s protocol. Libraries were packaged on dry ice and shipped overnight to Novogene Corporation Inc. (Sacramento, CA) to be sequenced using Illumina’s Hiseq 4000 for 150 bp paired-end sequencing. An average of 1,267,359 +/- 690,384 (σ) reads per sample were obtained.

### Taxonomic Assignment of Sequencing Data

For first sample set, 16S and ITS amplicon sequences were processed using Qiime2-2019.1.^24^ Demultiplexing was performed using the ‘q2-demux’ function with the ‘emp-paired’ preset. Sequencing reads were quality filtered, had chimeric sequences, PhiX, and singletons removed, and were clustered into amplicon sequence variants (ASVs) using the ‘q2-dada2’ function with the default parameters plus trunc_len_f = 280, trunc_len_r = 220, trim_left_f = 5, and trim_left_r = 5.^25^ This yielded 147 samples with an average of 30,051 +/- 24,768 (σ) high quality reads per sample for 16S amplicons (Supplemental Table 2), and 104 samples with an average of 3,517 +/-9,154 (σ) high quality reads for ITS amplicons (Supplemental Table 3). Taxonomic assignment of 16S and ITS reads was done using the ‘classify-sklearn’ function with default parameters. The databases used for classification were the Greengenes database (Version 13.8) for 16S data, and the UNITE database (Version 8.0) for ITS data.^26, 27^

For second sample set shotgun data, we first removed sequencing adapters using the ‘bbduk.sh’ script from BBMap v38.79 with the default parameters.^28^ Next, we demultiplexed our samples using ‘demuxbyname.sh’ script from BBMap using the default parameters. After demultiplexing, sequences were quality filtered using PRINSEQ++ v1.2 with the parameters trim_left = 5, trim_right = 5, min_len = 100, trim_qual_right = 28, and min_qual_mean = 25.^29^ This yielded an average of 1,209,001 +/- 643,544 (σ) high quality reads. Removal of human-derived reads was performed with Bowtie2 v2.3.5.1 on default settings by removing reads which aligned to the reference human genome, hg38.^30^ This final number of samples was 238, with an average of 1,102,247 +/- 643,325 (σ) high quality, non-human reads per sample (Supplemental Table 4). Lastly, we used IGGSearch v1.0 on the ‘lenient’ preset (--min-reads-gene=1 --min-perc-genes=15 --min-sp-quality=25) to characterize the taxonomy of our quality-filtered sequences.^31^

### Taxonomic Analysis

Data analysis was performed using R v3.6.3. For all sequencing runs, a synthetic microbial community DNA standard (ZymoBIOMICS D6305) was included as a control. When necessary, the first step in our compositional analysis was filtering taxa, from all samples, that contaminated the community standard control. Next, unassigned and mitochondrial reads were removed from our samples. 16S and ITS read counts were permutationally rarefied to 3,000 and 1,000 reads, respectively, for normalization purposes using the ‘rrarefy.perm’ function from the EcolUtils package v0.1. Shotgun data remained unrarefied. The alpha diversities for both amplicon and shotgun data were obtained using the ‘diversity’ and ‘specnumber’ functions from the Vegan v2.5-6 package, using the default parameters. Linear-mixed effect models (LME) were used for significance testing among alpha diversities to account for random effects, such as plate batching effects, and multiple measurements per individual using the nlme package, v3.1-148.

For all datasets, beta diversities were obtained using the ‘adonis’ function in Vegan to generate Bray-Curtis distance matrices and perform PERMANOVA significance testing from compositional data. PERMANOVA design and results can be found in Supplemental Tables 5-7 and Supplemental Tables 10-13. Significance was determined at p < 0.05 for both PERMANOVA and LME. Beta diversity was visualized using non-metric multidimensional scaling (NMDS) ordination obtained from the ‘metaMDS’ function in Vegan.

### Differential Abundance Testing

Our primary focus with the first sample set was to compare the microbial compositions of different sample types within the same individual. Therefore, we used ANCOM v2.1 in R to test for differentially abundant microbes since it can account for multiple variables and random effects.^32^ We used ANCOM with ‘sample type’ as our variable of interest (mucosal brushes vs. mucosal aspirates vs. colonoscopy lavage aspirates) and the individual of origin as a random effect. Other parameters included ‘p_adjust = FDR’ to control for the false discovery rate, and significance was determined at < 0.05.

For shotgun data, our primary focus was to compare the microbial composition of different subject types (Polyp-free vs. TA-bearing vs. SP-bearing). We used a univariate Kruskal-Wallis test with independent hypotheses weighting (IHW). IHW increases power while controlling the false discovery weight by utilizing covariate data that are independent of the null hypothesis.^33^ Before testing, we excluded samples with ‘Unknown/Other’ subject types, and filtered taxa that were not present in at least 20% of samples. We also eliminated repeated measurements by averaging the microbial relative abundances of left and right mucosal aspirates from the same individual. Kruskal-Wallis tests were performed for each taxon with the subject type as the variable. The IHW v1.14.0 package was used to correct p-values for the false discovery rate, using the sum of read counts per taxon across all samples as our covariate. FDR-adjusted p-values < 0.05 were considered significant. When visualizing relative abundances using a log_10_ scale, a pseudo-count of 0.0001 was added to prevent the removal of samples containing zeroes.

### Random Forests

Random Forests (RF) were performed on shotgun-sequenced mucosal aspirates to determine if the subject type of a sample could be predicted based on microbial composition. To do this, we used the rfPermute v1.9.3 package in R. We began by filtering taxa which were not present in at least 20% of mucosal aspirate samples. Two-thirds of the 156 shotgun mucosal aspirates were used for training the RF classifiers, while the remaining one-third was used for testing our RF models. RfPermute parameters were set to importance = TRUE, nrep = 100, ntree = 501, and mtry = 8. Afterwards, we generated receiver-operator curves (ROC) using the ‘roc’ function with default settings (pROC v1.18.0 package). Variables of importance were visualized with the ‘VarImpPlot’ function in the rfPermute package.

### Pathway Enrichment Analysis

Pathway enrichment analysis was done using unassembled shotgun reads with HUMAnN v3.0.1.^34^ The program was ran using the default parameters and the ChocoPhlAn v296 and UniRef90 v201901b databases were used for alignment. The ‘humann_renorm_table’ and ‘humann_join_tables’ functions were used to create a pathway abundance matrix of normalized counts in copies per million. Significantly enriched pathways between subject types were determined with a Kruskal-Wallis test using IHW. The false discovery rate was corrected for using the total sum of normalized counts per pathway as our covariate. Significance was determined at FDR < 0.05.

### Functional Metagenomic Analysis

Analysis of individual microbial genes was performed by cross-assembling reads into contiguous sequences using MEGAHIT v1.1.1.^35^ Contigs smaller than 2,500 bp were discarded and the remainder had open reading frames (ORFs) identified by Prodigal v2.6.3.^36^ The resulting ORFs were functionally annotated using eggNOG mapper v2.0, using the eggNOG v5.0 database. ^37^ Individual samples were aligned to annotated ORFs using Bowtie2 v2.3 to obtain per-sample ORF abundances. Per sample ORF abundances were compiled into a single ORF abundance table using the ‘pileup.sh’ script from BBMap. ORF counts were normalized to reads per kilobase per genome equivalent using MicrobeCensus v1.1.1 on default settings.^38^ Principal coordinate analysis was performed using the ‘cmdscale’ function from Vegan to visualize the functional metagenome composition among sample and subject types. PERMANOVA and differential abundance testing were performed in the same manner as with taxonomy.

## Results

### Microbiomes of Mucosal and Lavage Samples are similar to each other but different from those in Feces

To determine whether microbiome composition varied between sample types, we sequenced DNA from mucosal brushes, mucosal aspirates, and lavage aspirates from a subset of 38 individuals using 16S amplicon sequencing (Supplemental Table 2). We observed no significant differences in Shannon diversity or richness across mucosal brushes, mucosal aspirates, and lavage aliquots (Linear mixed effects model (LME): p > 0.05, Figure 2A). PERMANOVA analysis of Bray-Curtis dissimilarities revealed that the individual explained the greatest amount of variation in microbiome composition (R^2^ = 0.56, p = 0.001; Supplemental Table 5). This analysis found no significant differences in the microbiomes associated with mucosal brushes, mucosal aspirates, and lavage aliquots from within the same individual (R^2^ = 0.12, p = 0.49; Supplemental Table 5). The lack of significance was consistent with no discernable clusters based on sample type (Figure 2B). The abundances of only three ASV’s significantly differed across the three sampling methods – one from the *Gemellaceae* family and two *Streptococcus spp.* Abundances of these ASVs were higher in mucosal aspirates compared to mucosal brushes (ANCOM2: FDR < 0.05; Supplemental Figure 1).

**Figure 2:**
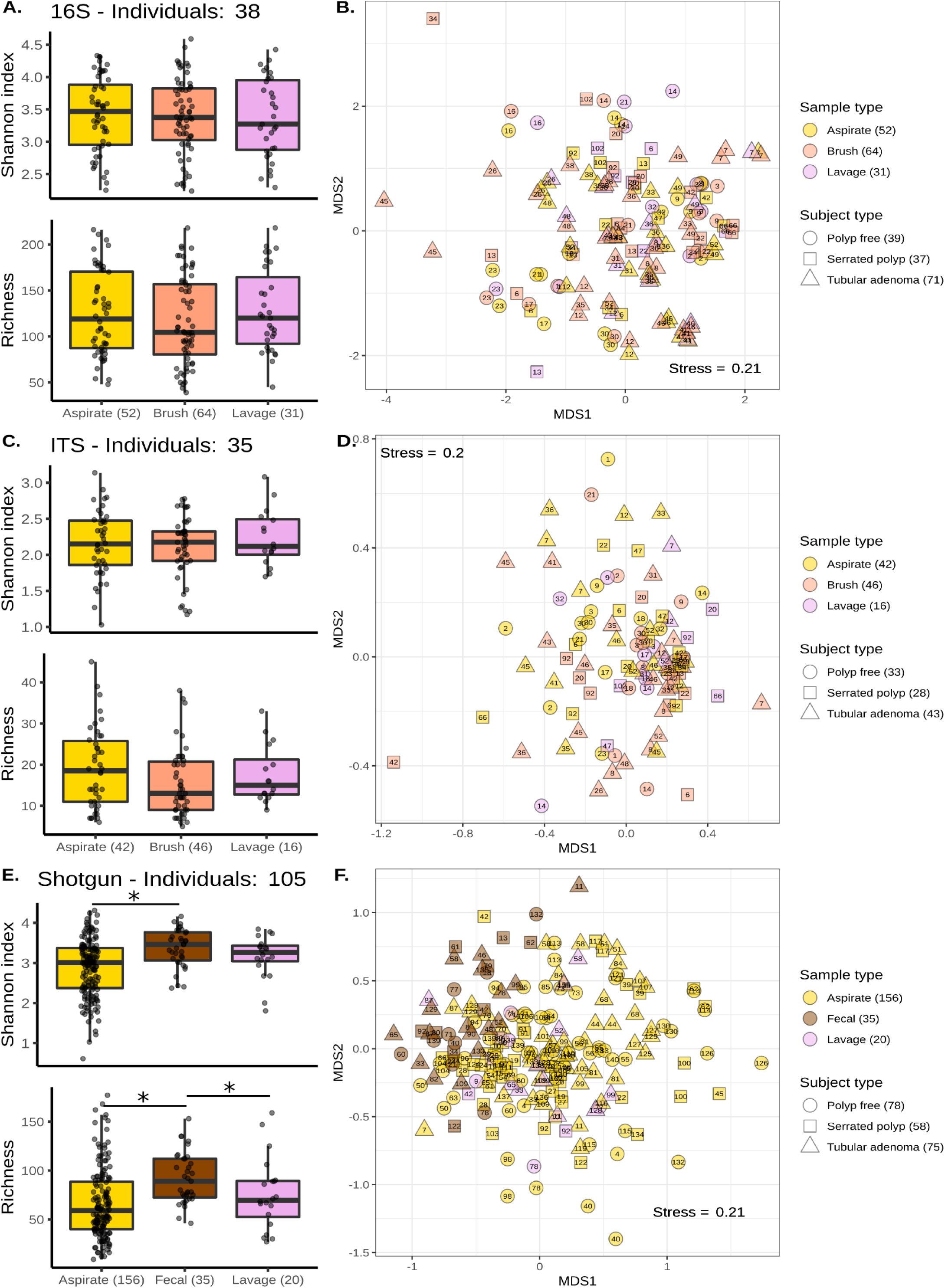
Microbiomes of Mucosal and Lavage Samples are similar to each other but different from those in Feces. **A, C, and E)** Box plots showing Shannon diversity and richness estimates across mucosal aspirates (yellow), mucosal brushes (orange), lavage aliquots (purple), and fecal samples (brown). The first sample set was sequenced using 16S **(A)**, and ITS **(C)** sequencing. The second sample set was sequenced using shotgun sequencing **(E)**. The center line within each box defines the median, boxes define the upper and lower quartiles, and whiskers define 1.5x the interquartile range. **B, D, and F)** Non-metric multidimensional scaling of Bray-Curtis dissimilarities produced from 16S **(B)**, ITS **(D)**, and shotgun **(F)** compositional data. Each point corresponds to one sample, with multiple samples per individual. The individual of origin is denoted numerically within each point. The number of samples per sample type and subject category are annotated parenthetically. Significant comparisons (p < 0.05) are denoted by an asterisk (*).

ITS2 sequencing was also performed on the same subset of samples to investigate the effect of sampling method on the fungal microbiome (Supplemental Table 3). We observed no differences in Shannon diversity or richness across mucosal brushes, mucosal aspirates, and lavage aliquots (LME: p > 0.05, Figure 2C). Beta-diversity ordination by sample type demonstrated no discernable clustering (Figure 2D). Like 16S amplicon data, PERMANOVA analysis of Bray-Curtis dissimilarities showed that the individual explained the greatest amount of variation in fungal community composition (R^2^ = 0.28, p < 0.001), with no significant associations between fungal community composition and our three sampling methods (R^2^ = 0.38, p = 0.123; Supplemental Table 6).

Following the collection of fecal samples, we performed shotgun sequencing on a second subset of samples, representing 117 individuals (Supplemental Table 4). Mucosal brushes were excluded from the second sample set because a pilot shotgun sequencing run revealed these samples contained a large percentage of human-derived reads (Supplemental Figure 2). Based on estimates of Shannon diversity and species richness, the microbiomes in fecal samples were significantly more diverse than those in the mucosal aspirates (LME: p = 0.007 and p = 0.002, respectively) and marginally more diverse than those in lavage aliquots (LME: p = 0.053 and p = 0.047, respectively; Figure 2E). Visualization of sample beta-diversities revealed a cluster of fecal samples that partially overlapped with mucosal and lavage aspirates (Figure 2F). PERMANOVA showed that the individual explained the greatest amount of variation in microbiome composition (R^2^ = 0.75, p < 0.001; Supplemental Table 7). In comparison, sampling method explained 14% of variation in the microbiome (PERMANOVA: p = 0.001). Fecal samples had a mean relative abundance of 63% for Firmicutes, 27% for Bacteroides, 3.5% for Actinobacteria, and 4.5% for Proteobacteria. Mucosal aspirates and lavage aliquots were more similar and had a mean relative abundance of 73% and 75% for Firmicutes, 15% and 11% for Bacteroides, 4.7% and 5.2% for Actinobacteria, and 4.0% and 6.6% for Proteobacteria, respectively (Supplemental Figure 3). Differential abundance analysis revealed 44 OTUs whose abundances significantly differed between fecal samples and mucosal aspirates (ANCOM2: FDR < 0.05; Supplemental Table 8). Six OTUs were differentially abundant between fecal samples and lavage aliquots (Supplemental Table 9), and no OTUs were significantly different between mucosal aspirates and lavage aliquots (ANCOM2; FDR > 0.05).

### The Microbiomes of Polyps and Opposite Wall Healthy Tissue are similar within Individuals

To identify potential polyp-specific microbial biomarkers, 14 mucosal brush samples from 6 individuals were collected from polyps and opposite wall healthy tissue and sequenced as part of the first sample set (Figure 3A). Based on 16S sequencing, we observed no significant differences in Shannon diversity or richness between polyp and opposite wall healthy tissue from within the same individual (Figure 3B). Similarly, there were no significant differences in beta-diversity across polyp and opposite wall healthy tissue pairs (PERMANOVA: R^2^ = 0.19, p = 0.53; Figure 3C; Supplemental Table 10). We were unable to identify any differentially abundant microbes between polyp and opposite wall tissue brushes. Microbiomes were mostly individualistic, with subject origin explaining 52% of the variance in microbiome composition (PERMANOVA: p = 0.004; Figure 3D; Supplemental Table 10). We detected significant associations between microbiome composition and colon side (right/proximal versus left/distal), representing 16% of the observed variance (PERMANOVA: p = 0.005; Supplemental Table 10). Significant associations within the microbiome were observed when both polyp and opposite wall tissue pairs were categorized by subject type, explaining approximately 10% of variance (PERMANOVA: p = 0.03; Supplemental Table 10).

**Figure 3:**
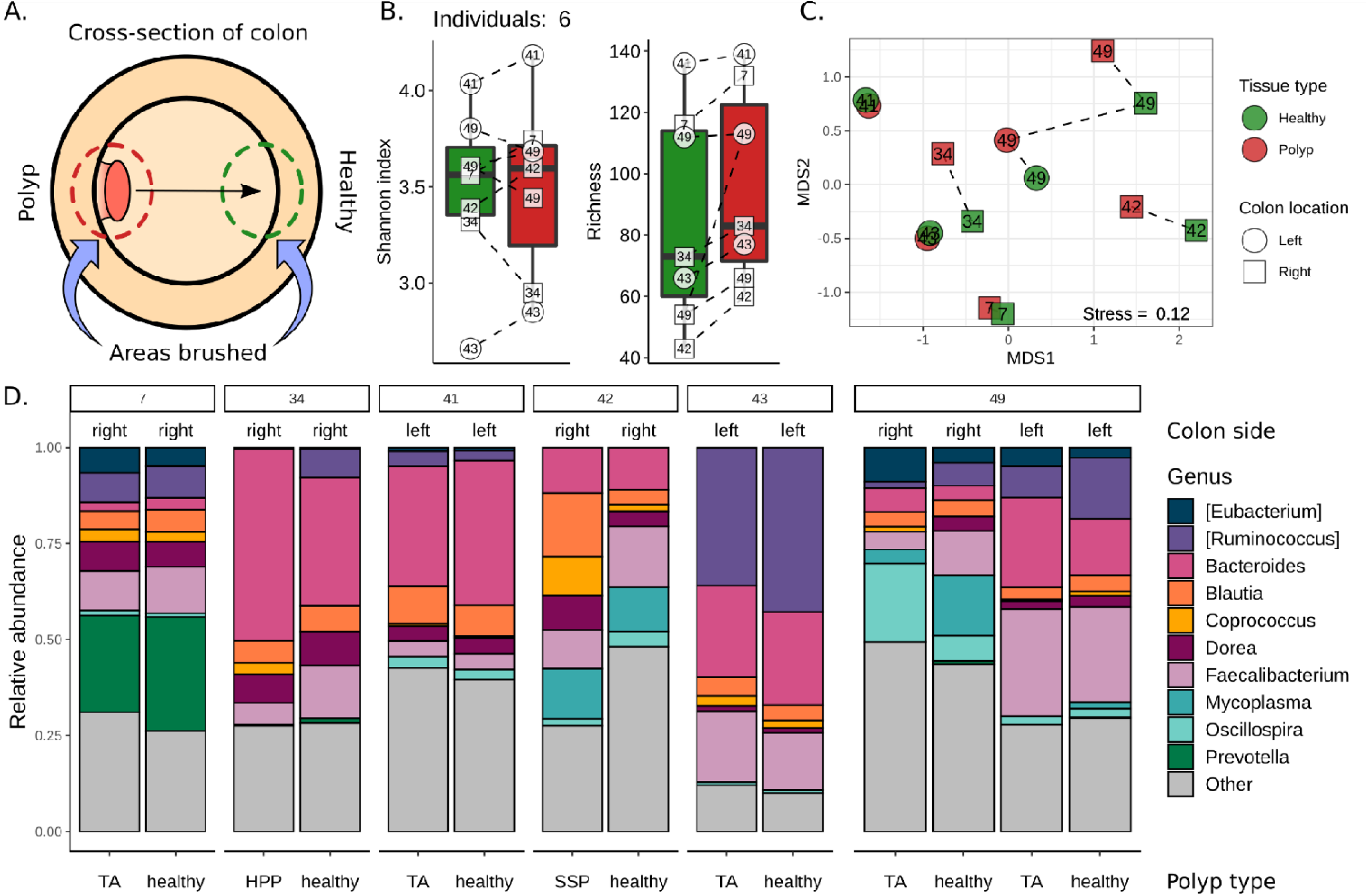
The Microbiomes of Polyps and Opposite Wall Healthy Tissue are similar within Individuals. **A)** An illustration of the sampling strategy used to characterize the microbial community of 16S mucosal brushes from polyps (red) and opposite wall tissue (green). **B)** Box plots of Shannon diversity and richness estimates from polyp and opposite wall brushes. The center line within each box defines the median, boxes define the upper and lower quartiles, and whiskers define 1.5x the interquartile range. **C)** Non-metric multidimensional scaling of Bray-Curtis dissimilarities of polyp and opposite wall tissue brushes. Each point is one sample, with multiple samples per individual. The individual of origin is denoted numerically within each point. The shape of each point denotes the right (proximal) and left (distal) side of the colon. **D)** The relative abundance of the top ten microbial genera across all samples. Samples are grouped by each individual and labeled by polyp type, where tubular adenoma = TA, hyperplastic polyp = HPP, and sessile serrated polyp = SSP.

### Tubular Adenoma-bearing, Serrated Polyp-bearing, and Healthy Individuals have distinct Microbiomes

We next reanalyzed all samples from the first and second sample sets to examine whether the subject type of a sample (polyp-free, TA-bearing, or SP-bearing) was significantly associated with microbial diversity and composition. In both 16S and shotgun data, we observed no significant differences between subject types based on Shannon diversity or richness estimates (LME: p > 0.05; Supplemental Figure 4). In ITS data, we observed significantly increased Shannon diversity, but not richness, in samples from polyp-free individuals when compared to those from TA-bearing individuals (LME: p = 0.03; Supplemental Figure 4). Beta diversity analysis of 16S and ITS data from the first sample set demonstrated that subject type explained 5% and 2% of the variance associated with the microbiome, respectively (16S PERMANOVA: p = 0.001; Supplemental Table 5 and ITS PERMANOVA: p = 0.11; Supplemental Table 6). A similar result was observed in the second sample set. We found significant associations between the microbiome and subject type, explaining 2% of the variance observed (PERMANOVA: p = 0.001; Supplemental Table 7). This association between microbiome composition and subject type was not observed when testing lavage aliquots (PERMANOVA: p = 0.47; Supplemental Table 11) or fecal samples (PERMANOVA: p = 0.10; Supplement table 12) alone. Taxonomic visualization suggested that TA-bearing mucosal aspirates were distinct compared to polyp-free and SP-bearing mucosal aspirates (Figure 4A).

**Figure 4:**
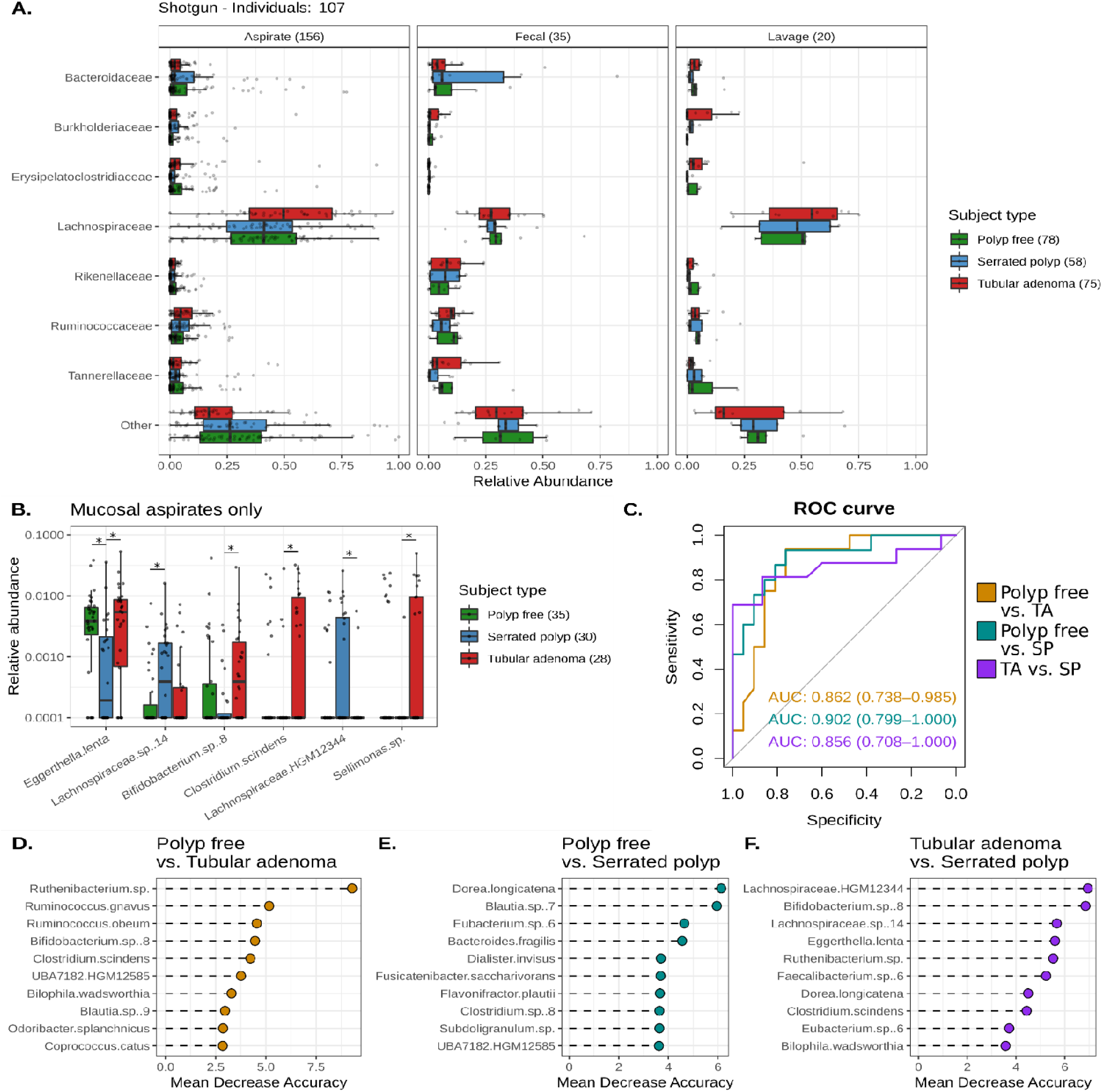
Tubular Adenoma-bearing, Serrated Polyp-bearing, and Healthy Individuals have distinct Microbiomes. **A)** Box plots of the top seven most abundant microbial families across all samples from the second sample set. The number of samples per sampling method and subject type are denoted parenthetically, with multiple samples per individual. **B)** Box plots showing the relative abundances of microbes determined to be differentially abundant between each subject type using shotgun-sequenced mucosal aspirates. Each point refers to a single individual. Significant comparisons (p < 0.05) are denoted by an asterisk (*). The center line within each box defines the median, boxes define the upper and lower quartiles, and whiskers define 1.5x the interquartile range. **C)** A receiver operating characteristic (ROC) curve illustrating the true positive rate (Sensitivity, y-axis) versus the false positive rate (Specificity, x-axis) produced by Random Forest classification. The area under the curve (AUC) value for each Random Forest is displayed with the 90% confidence interval. **D)** The top ten variables of importance for each pairwise random forest classification. Variables are sorted by their mean decrease in accuracy, with larger means contributing greater to random forest performance.

We next performed an in-depth investigation of each subject type’s microbiome using second sample set mucosal aspirates, due to their larger comparable sample size. Differential abundance analysis demonstrated that six microbes were significantly different in at least one subject type comparison (Figure 4B). *Eggerthella lenta* was the most significantly different taxon, and it was depleted in SP-bearing samples when compared to polyp-free (FDR = 3 x 10^-3^)and TA-bearing (FDR = 9 x 10^-3^) samples. *Bifidobacterium* sp. was also depleted in SP-bearing samples but the difference was only significant when compared to TA-bearing samples (FDR = 0.01). Conversely, two species of *Lachnospiraceae* were enriched in SP-bearing samples when compared to polyp-free (*Lachnospiraceae* sp. FDR = 0.04) and TA-bearing samples (*Lachnospiraceae* HGM12344 FDR = 0.02). *Clostridium scindens* and *Sellimonas* sp. were both enriched in TA-bearing mucosal aspirates when compared SP-bearing aspirates (FDR = 8 x 10^-3^ for *C. scindens,* FDR = 4 x 10^-3^ for *Sellimonas* sp.). Supplemental Figure 5 suggest that *E. lenta* was also depleted in 16S mucosal aspirates, but the result was not statistically significant.

We next examined whether microbial composition could predict the subject type of mucosal aspirates. Random forest (RF) was able to accurately classify mucosal aspirates from each pairwise subject type comparison, producing area under curve (AUC) values of at least 85% for each comparison (Figure 4C). The microbes most important for determining the classification of polyp-free versus TA-bearing mucosal aspirates were *Ruthenibacterium* sp., *Ruminococcus gnavus, Ruminococcus obeum,* and the previously observed *Bifidobacterium* sp. and *C. scindens* (Figure 4D). For the polyp-free versus SP-bearing RF classification, *Dorea longicatena, Blautia* sp., *Eubacterium* sp., and *Bacteroides fragilis* were the most important variables (Figure 4E). Lastly, *Lachnospiraceae* HGM12344, *Bifidobacterium* sp., *Lachnospiraceae* sp., and *E. lenta* were the top microbes of importance for the SP-bearing versus TA-bearing RF classification (Figure 4F). Supplemental Figure 6 displays the relative abundances of the top variables of importance for all RF comparisons. These data suggest each subject type has a distinct microbial composition which can be used to predict the origin of mucosal aspirates.

### Microbiome Functional Potential is distinct across Sampling Methods and Subject Types

The functional characteristics of our shotgun metagenomes were next explored. Pathway analysis was performed, which resulted in the discovery of 507 metabolic pathways (Supplemental Figure 7). Of those pathways, only the 1,5-anhydrofructose degradation pathway was significantly more abundant in TA-bearing mucosal aspirates when compared to polyp-free ones. (FDR = 0.03, Figure 5A). Subsequently, we analyzed individually annotated microbial genes. Principal coordinate analysis produced a result similar to our taxonomic ordination, with fecal samples clustering together and no obvious subject type clustering (Figure 5B). This was supported by PERMANOVA, which confirmed an association between functional metagenome and sampling method, explaining 10.9% of the observed variance (PERMANOVA: p = 0.001; Supplemental Table 13). By comparison, the individual of origin explained approximately 75% of the observed variance in the functional microbiome (PERMANOVA: p = 0.001; Supplemental Table 13) and subject type explained 1.6% of the observed variance (PERMANOVA: p = 0.001; Supplemental Table 13).

**Figure 5:**
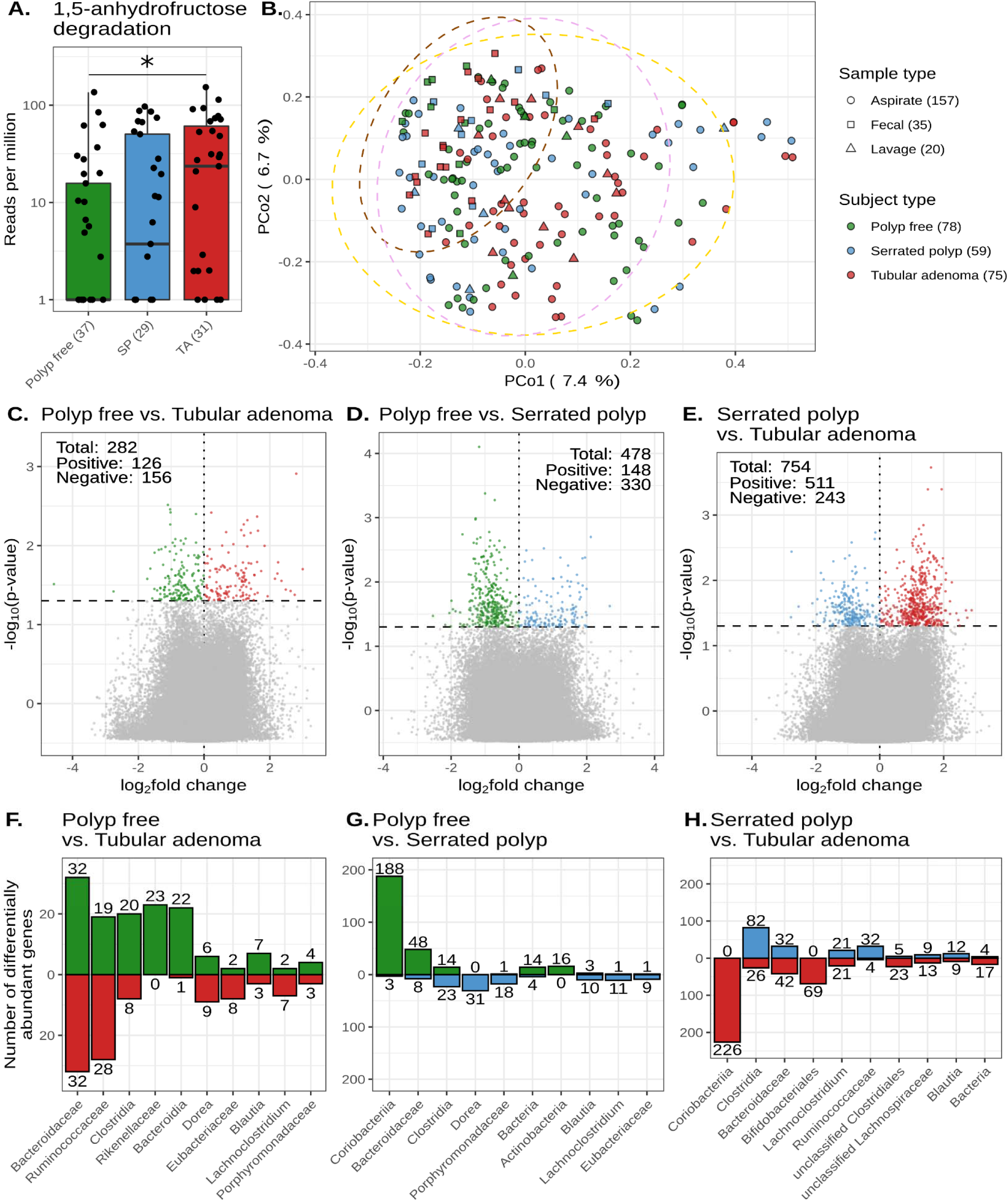
Microbiome Functional Potential is distinct across Subject Types. **A)** A box plot displaying the abundance in reads-per-million of the 1,5-anhydrofructose degradation pathway in mucosal aspirates. Each point represents a single individual and significant comparisons are denoted with an asterisk (*). The center line within each box defines the median, boxes define the upper and lower quartiles, and whiskers define 1.5x the interquartile range. **B)** Principal coordinate analysis of per-gene Bray-Curtis dissimilarities across second sample set mucosal aspirates, lavage aliquots, and fecal samples. Ellipses are drawn to represent the 95% confidence interval of each sample type’s distribution. Points represent a single sample, with multiple samples per individual. The number of samples per sampling method and subject type are annotated parenthetically. **C, D, and E)** Volcano plots illustrating the differential abundances of microbial genes in mucosal aspirate samples. The horizontal and vertical lines denote the significance threshold of p = 0.05, and a log2fold change of zero, respectively. Points are colored to denote the subject type in which the gene was more abundant, with green referring to genes more abundant in polyp-free samples, red for tubular adenomas, and blue for serrated polyps. The number of total, negative fold-change, and positive-fold change genes are displayed within each graph. **F, G, and H)** The number of differentially abundant genes per taxon for each subject type comparison. Only the top ten taxa with the most differentially abundant genes are shown.

We concluded our analysis by examining the differentially abundant genes among subject types using mucosal aspirates. There were 282 differentially abundant genes between polyp-free and TA-bearing mucosal aspirates (Figure 5C). Of those genes, 126 had a positive log_2_ fold change (more abundant in TA-bearing), and 156 had a negative log_2_ fold change (more abundant in polyp-free). Figure 5F displays the most resolved taxonomic level for the top ten taxa with the most differentially abundant genes. For polyp-free and TA-bearing samples, the *Bacteroidaceae* family contained 64 of the 282 differentially abundant genes. Comparatively, 478 differentially abundant genes were present between polyp-free and SP-bearing mucosal aspirates (Figure 5D). These genes were mostly more abundant in polyp-free samples, with 330 genes having a negative log_2_ fold change and 148 having positive log_2_ fold change. *Coriobacteriia*, the class to which *E. lenta* belongs, was responsible for 188 of the 478 differentially abundant genes (Figure 5G). A similar result was observed when comparing SP-bearing to TA-bearing samples, as of the 754 differentially abundant genes, 226 belonged to *Coriobacteriia* (Figure 5E and 5H). A complete list of all the differentially abundant genes, their functions, and taxonomy can be found in the supplement (Supplemental File 1).

## Discussion

### Sampling Method and Microbiome Characterization

In this study we used direct and indirect methods to sample the colon and characterize the microbiomes of polyp-free and colorectal polyp-bearing individuals. Using amplicon sequencing, we found that microbiomes of mucosal brushes and mucosal aspirates did not significantly differ in diversity or composition. In contrast, the microbiomes of fecal samples were significantly more diverse and compositionally distinct when compared to those from mucosal aspirates.

Due to their ease of collection, fecal samples are frequently used to study the human microbiome in the context of CRC. However, fecal samples poorly represent the microbiota adherent to the colon mucosa, and instead capture those found in the intestinal lumen.^20, 21^ Their increased diversity and paucity of mucosa-associated microbes suggests that fecal samples are not ideal for discovering novel CRC biomarkers. This is especially true for premalignant polyps, which have less pronounced signatures of microbial dysbiosis when compared to carcinomas. This is supported by Peters *et al.*, who found more pronounced compositional changes in the microbiomes of fecal samples from advanced conventional adenomas when compared to those from non-advanced adenomas.^17^ The decreased sensitivity of fecal samples to detect CRC-associated microbes is also highlighted by their results demonstrating significant associations between the gut microbiome and distal conventional adenoma cases, but not proximal.^17^ This is also likely why Peters *et al.* did not observe substantial differences in the microbial compositions of HPP, SSP, and healthy samples, as serrated polyps predominantly develop in the proximal colon.

In this study, we reported significant associations between the gut microbiome and mucosal aspirates obtained from both the proximal and distal colon. We also observed significant differences when comparing the microbiomes of polyp-free samples to SP-bearing ones using mucosal aspirates. No such differences were seen in fecal samples, but this may be a result of smaller sample size. Nevertheless, these data suggest that mucosal samples are sensitive enough to study the microbiome of colorectal polyps found within the proximal colon. These results contradict a study published by Yoon *et al.*, who did not find significant compositional differences the in mucosa-associated gut microbiota among polyp-free, TA, SSP, and CRC bearing individuals.^18^ The authors did note, however, that this result was likely driven by the small sample size of the study, with only 6 samples per group, and 24 samples total.

Compared to mucosal brushes, mucosal aspirates had a lower risk of damaging the host epithelium, provided larger collection volumes for downstream sample processing, and resulted in a lower proportion of human derived reads during shotgun sequencing. Because of these advantages, we recommend using mucosal aspirates rather than mucosal brushing for characterizing the microbiomes of colorectal polyps.

### Hyperlocal Microbiome Comparisons

Direct sampling of polyp mucosa with brushes revealed no differences in the hyperlocal microbiome of polyp tissue versus opposite colon wall tissue. One factor which could have disrupted the potential hyperlocal differences in the gut microbiota is the colonoscopy preparation and lavage. As part of the preparation, individuals were advised to adhere to a low fiber, clear liquid diet 24 hours prior to colonoscopy. Dietary fiber is important in maintaining the longitudinal and lateral organization of the gut microbiota within the colon, as mice on a low fiber diet show disrupted microbial organization.^20^ Changes in diet can rapidly shift the composition of the gut microbiome, often within 24 hours, in both humans and mice.^7, 39, 40^ A laxative-based cleansing and colonoscopy rinse was also performed, potentially obscuring the hyperlocal organization further. Nevertheless, significant compositional differences between the microbiomes of samples taken from the proximal and distal colon were observed, suggesting that broad microbial organization remained present in the gut after colonoscopy preparation and lavage. It is important to note that these claims are based on data from 14 samples from 6 individuals, therefore, additional studies with more samples are needed to validate the reproducibility of our findings.

### Microbiome Signatures of the CRC Carcinogenesis Pathways

Compositional differences were observed in the gut microbiome across TA-bearing, SP-bearing, and polyp-free individuals using mucosal sampling. Notably, we demonstrated that the microbial composition of each subject type was distinct enough to accurately predict the origin of mucosal aspirates using RF. This is further evidenced by the discovery of 6 and 1,143 differentially abundant taxa and genes among subject types, respectively. These findings suggest that the gut microbiome functions differently between the adenoma-carcinoma sequence and the serrated pathway.

In the adenoma-carcinoma sequence, the gut microbiome exists in, and potentially contributes to, an inflammatory environment to promote colorectal carcinogenesis. Data obtained from mucosal aspirates also support this theory. RF classification identified *Ruminococcus gnavus* and *Bacteroides fragilis* as top variables of importance, both of which were elevated in TA-bearing mucosal aspirates. *R. gnavus* has been previously associated with CRC and inflammatory bowel disease.^41^ *B. fragilis* produces a metalloprotease that causes oxidative DNA damage and cleaves the tumor suppressor protein, E-cadherin.^42–44^ *C. scindens* was also significantly elevated in TA-bearing aspirates, and it can metabolize excess primary bile acids not absorbed by the small intestine into secondary bile acids.^45, 46^ We did not observe an increased abundance of *C. scindens* bile acid metabolism genes directly, but high concentrations of secondary bile acids can cause host oxidative stress, nitrosative stress, DNA damage, apoptosis, and mutations.^47^ Secondary bile acids also act as farnesoid X receptor antagonists, resulting in enhanced *wnt* signaling in the adenoma-carcinoma sequence.^48^ High fructose corn syrup consumption has also been associated with increased CRC risk.^49, 50^ Here, we observed that the TA-associated microbiome of mucosal aspirates had an increased abundance of genes that encode the pathway for degrading 1,5-anhydrofructose, which can be produced from fructose, glucose, or starch.^51^ This pathway was elevated in TA-bearing samples, but 1,5-anhydrofructose has been shown to promote the growth of the beneficial microbe, *Faecalibacterium prausnitzii*, and reduce inflammation by inactivating NF-κB.^52, 53^ *Sellimonas*, which has been found to negatively correlate with clinical tumor markers, was more also abundant in TA-bearing mucosal aspirates.^54^

Unlike the adenoma-carcinoma sequence, the microbiome in the serrated pathway remains understudied. *Fusobacterium nucleatum,* which has been implicated in the adenoma-carcinoma sequence because of its ability to activate *wnt* signaling, has also been described as having a role in serrated CRC development.^55^ *F. nucleatum* abundance is associated with serrated pathway lesions and features, such as mismatch repair deficiency, MLH1 methylation, CpG island methylator phenotype, and high microsatellite instability.^14^ Here, we did not find differences in *F. nucleatum* abundances across HPPs, SSPs, TAs, or polyp-free controls. Instead, we most prominently found that *E. lenta* was depleted in mucosal aspirates from SP-bearing individuals, a result that spanned 16S and shotgun data.

*E. lenta* metabolizes inert plant lignans in the gut into bioactive enterolignans, such as enterolactone and enterodiol.^56^ These enterolignans have anti-proliferative and anti-inflammatory effects, and help modulate estrogen signaling, lipid metabolism, and bile acid regulation.^57^ They have also been associated with reduced cancer risk.^58^ Diets rich in plant fiber have been associated with decreased CRC risk.^6, 59^ Fiber is fermented by the intestinal microbiota to produce short chain fatty acids, including acetate, butyrate, and propionate. Butyrate is the primary energy source for colonocytes and has anti-inflammatory and anti-tumor properties.^60–62^ Butyrate also is involved in the epigenetic expression of genes as a histone deacetylase inhibitor.^63^ In the serrated pathway, the gene SLC5A8, which mediates short chain fatty acid uptake into colonic epithelial cells, is frequently inhibited via promoter methylation, suggesting that dietary fiber may be required for proper cellular epigenetic regulation.^64^

More evidence of dietary fiber playing a role in the SP-associated microbiome is the depletion of many carbohydrate active enzymes in our functional metagenomic data. This includes a 4-amino-4-deoxy-L-arabinose transferase and a glycosyltransferase family 2 from *E. lenta*, *axeA* from *Bifidobacterium,* and *uidA, sacC,* and a glycoside hydrolase family 31 from the *Bacteroidaceae*. Therefore, we hypothesize that low dietary fiber consumption facilitates aberrant epigenetic modifications within colonocytes to promote serrated polyp development. Studies which utilize both mucosal sampling methods and dietary information are needed to test this hypothesis.

## Conclusion

The complex and individualistic nature of the human gut microbiome has made it difficult to mechanistically link the microbiome with colorectal carcinogenesis. By describing the association between the gut microbiota and serrated polyps, our study provides novel insight into potential mechanisms for the epigenetic-based serrated pathway of CRC. In addition, our data underscores the importance of distinguishing between different pathways of colorectal carcinogenesis when investigating the gut microbiome. Finally, transitioning future microbiome studies to mucosal sampling methods will enable the discovery of previously unassociated microbes as demonstrated here.

## Declarations

### Author contributions

Katrine L. Whiteson and William E. Karnes devised the study design with support from Lauren DeDecker. Subject recruitment was performed by William E. Karnes and Zachary Lu. Sample collection was performed by William E. Karnes, Lauren DeDecker, and Zachary Lu with guidance from Katrine L. Whiteson. Julio Avelar-Barragan, Bretton Coppedge, and Zachary Lu processed samples for data acquisition. Julio Avelar-Barragan performed the data analysis and wrote the manuscript with guidance from Katrine L. Whiteson.

### Ethics approval

This study was approved by the Institutional Review Board (IRB) of the University of California, Irvine (HS# 2017-3869).

### Data availability statement

All code for data processing and analysis is available on GitHub at: https://github.com/Javelarb/ACS_polyp_study. Additional data and materials are available upon reasonable request.

### Data deposition

Sequencing data is available on the Sequence Read Archive under the BioProject ID, PRJNA745329.

## Acknowledgements

We would like to thank Claudia Weihe and Jennifer B.H Martiny for allowing us to borrow laboratory equipment and giving insightful feedback, Andrew Oliver and Jason A. Rothman for their bioinformatic expertise, Clark Hendrickson for his assistance with sample preparation, and Heather Maughan for her helpful edits and suggestions.

## Abbreviations

CRC: Colorectal cancer
APC: Adenomatous polyposis coli
HPP: Hyperplastic polyp
TSA: Traditional serrated adenoma
SSP: Sessile serrated polyp
SP: Serrated polyp
TA: Tubular adenoma
OTU: Operational taxonomic unit
ASV: Amplicon sequence variant
ORF: Open reading frame
LME: Linear mixed effects model
RF: Random Forest

## REFERENCES

1. Sung, Hyuna, Jacques Ferlay, Rebecca L. Siegel, Mathieu Laversanne, Isabelle Soerjomataram, Ahmedin Jemal, and Freddie Bray. “Global Cancer Statistics 2020: GLOBOCAN Estimates of Incidence and Mortality Worldwide for 36 Cancers in 185 Countries.” CA: A Cancer Journal for Clinicians 71, no. 3 (May 2021): 209–49. https://doi.org/10.3322/caac.21660.

2. Stoffel, Elena M., Pamela B. Mangu, Stephen B. Gruber, Stanley R. Hamilton, Matthew F. Kalady, Michelle Wan Yee Lau, Karen H. Lu, Nancy Roach, and Paul J. Limburg. “Hereditary Colorectal Cancer Syndromes: American Society of Clinical Oncology Clinical Practice Guideline Endorsement of the Familial Risk–Colorectal Cancer: European Society for Medical Oncology Clinical Practice Guidelines.” Journal of Clinical Oncology 33, no. 2 (January 10, 2015): 209–17. https://doi.org/10.1200/JCO.2014.58.1322.

3. Collins, S.M., E. Denou, E.F. Verdu, and P. Bercik. “The Putative Role of the Intestinal Microbiota in the Irritable Bowel Syndrome.” Digestive and Liver Disease 41, no. 12 (December 2009): 850–53. https://doi.org/10.1016/j.dld.2009.07.023.

4. Verdam, Froukje J., Susana Fuentes, Charlotte de Jonge, Erwin G. Zoetendal, Runi Erbil, Jan Willem Greve, Wim A. Buurman, Willem M. de Vos, and Sander S. Rensen. “Human Intestinal Microbiota Composition Is Associated with Local and Systemic Inflammation in Obesity: Obese Gut Microbiota and Inflammation.” Obesity 21, no. 12 (December 2013): E607–15. https://doi.org/10.1002/oby.20466.

5. Song, Mingyang, Wendy S. Garrett, and Andrew T. Chan. “Nutrients, Foods, and Colorectal Cancer Prevention.” Gastroenterology 148, no. 6 (May 2015): 1244–1260.e16. https://doi.org/10.1053/j.gastro.2014.12.035.

6. “Diet, Nutrition, Physical Activity, and Colorectal Cancer.” World Cancer Research Fund/American Institute for Cancer Research. Continuous Update Project Expert Report, 2018. dietandcancerreport.org.

7. David, Lawrence A., Corinne F. Maurice, Rachel N. Carmody, David B. Gootenberg, Julie E. Button, Benjamin E. Wolfe, Alisha V. Ling, et al. “Diet Rapidly and Reproducibly Alters the Human Gut Microbiome.” Nature 505, no. 7484 (January 2014): 559–63. https://doi.org/10.1038/nature12820.

8. Engen, Phillip A., Stefan J. Green, Robin M. Voigt, Christopher B. Forsyth, and Ali Keshavarzian. “The Gastrointestinal Microbiome: Alcohol Effects on the Composition of Intestinal Microbiota.” Alcohol Research: Current Reviews 37, no. 2 (2015): 223–36.

9. Ley, Ruth E. “Obesity and the Human Microbiome:” Current Opinion in Gastroenterology 26, no. 1 (January 2010): 5–11. https://doi.org/10.1097/MOG.0b013e328333d751.

10. Mailing, Lucy J., Jacob M. Allen, Thomas W. Buford, Christopher J. Fields, and Jeffrey A. Woods. “Exercise and the Gut Microbiome: A Review of the Evidence, Potential Mechanisms, and Implications for Human Health.” Exercise and Sport Sciences Reviews 47, no. 2 (April 2019): 75–85. https://doi.org/10.1249/JES.0000000000000183.

11. Nakanishi, Yuki, Maria T. Diaz-Meco, and Jorge Moscat. “Serrated Colorectal Cancer: The Road Less Travelled?” Trends in Cancer 5, no. 11 (November 2019): 742–54. https://doi.org/10.1016/j.trecan.2019.09.004.

12. Pino, Maria S., and Daniel C. Chung. “The Chromosomal Instability Pathway in Colon Cancer.” Gastroenterology 138, no. 6 (May 2010): 2059–72. https://doi.org/10.1053/j.gastro.2009.12.065.

13. De Palma, Fatima, Valeria D’Argenio, Jonathan Pol, Guido Kroemer, Maria Maiuri, and Francesco Salvatore. “The Molecular Hallmarks of the Serrated Pathway in Colorectal Cancer.” Cancers 11, no. 7 (July 20, 2019): 1017. https://doi.org/10.3390/cancers11071017.

14. DeDecker, Lauren, Bretton Coppedge, Julio Avelar-Barragan, William Karnes, and Katrine Whiteson. “Microbiome Distinctions between the CRC Carcinogenic Pathways.” Gut Microbes, January 15, 2021, 1–12. https://doi.org/10.1080/19490976.2020.1854641.

15. Kahi, Charles J. “Screening Relevance of Sessile Serrated Polyps.” Clinical Endoscopy 52, no. 3 (May 31, 2019): 235–38. https://doi.org/10.5946/ce.2018.112.

16. Delker, Don A., Brett M. McGettigan, Priyanka Kanth, Stelian Pop, Deborah W. Neklason, Mary P. Bronner, Randall W. Burt, and Curt H. Hagedorn. “RNA Sequencing of Sessile Serrated Colon Polyps Identifies Differentially Expressed Genes and Immunohistochemical Markers.” Edited by Frank T. Kolligs. PLoS ONE 9, no. 2 (February 12, 2014): e88367. https://doi.org/10.1371/journal.pone.0088367.

17. Peters, Brandilyn A., Christine Dominianni, Jean A. Shapiro, Timothy R. Church, Jing Wu, George Miller, Elizabeth Yuen, et al. “The Gut Microbiota in Conventional and Serrated Precursors of Colorectal Cancer.” Microbiome 4, no. 1 (December 2016): 69. https://doi.org/10.1186/s40168-016-0218-6.

18. Yoon, Hyuk, Nayoung Kim, Ji Hyun Park, Yong Sung Kim, Jongchan Lee, Hyoung Woo Kim, Yoon Jin Choi, et al. “Comparisons of Gut Microbiota Among Healthy Control, Patients With Conventional Adenoma, Sessile Serrated Adenoma, and Colorectal Cancer.” Journal of Cancer Prevention 22, no. 2 (June 30, 2017): 108–14. https://doi.org/10.15430/JCP.2017.22.2.108.

19. Rezasoltani, Sama, Hamid Asadzadeh Aghdaei, Hossein Dabiri, Abbas Akhavan Sepahi, Mohammad Hossein Modarressi, and Ehsan Nazemalhosseini Mojarad. “The Association between Fecal Microbiota and Different Types of Colorectal Polyp as Precursors of Colorectal Cancer.” Microbial Pathogenesis 124 (November 2018): 244–49. https://doi.org/10.1016/j.micpath.2018.08.035.

20. Riva, Alessandra, Orest Kuzyk, Erica Forsberg, Gary Siuzdak, Carina Pfann, Craig Herbold, Holger Daims, Alexander Loy, Benedikt Warth, and David Berry. “A Fiber-Deprived Diet Disturbs the Fine-Scale Spatial Architecture of the Murine Colon Microbiome.” Nature Communications 10, no. 1 (December 2019): 4366. https://doi.org/10.1038/s41467-019-12413-0.

21. Chen, Weiguang, Fanlong Liu, Zongxin Ling, Xiaojuan Tong, and Charlie Xiang. “Human Intestinal Lumen and Mucosa-Associated Microbiota in Patients with Colorectal Cancer.” Edited by Antonio Moschetta. PLoS ONE 7, no. 6 (June 28, 2012): e39743. https://doi.org/10.1371/journal.pone.0039743.

22. Looby, Caitlin I., Mia R. Maltz, and Kathleen K. Treseder. “Belowground Responses to Elevation in a Changing Cloud Forest.” Ecology and Evolution 6, no. 7 (April 2016): 1996– 2009. https://doi.org/10.1002/ece3.2025.

23. Weihe, Claudia, and Avelar-Barragan, Julio. “Next Generation Shotgun Library Preparation for Illumina Sequencing - Low Volume V1.” Accessed January 3, 2022. https://doi.org/10.17504/protocols.io.bvv8n69w.

24. Bolyen, Evan, Jai Ram Rideout, Matthew R. Dillon, Nicholas A. Bokulich, Christian C. Abnet, Gabriel A. Al-Ghalith, Harriet Alexander, et al. “Reproducible, Interactive, Scalable and Extensible Microbiome Data Science Using QIIME 2.” Nature Biotechnology 37, no. 8 (August 2019): 852–57. https://doi.org/10.1038/s41587-019-0209-9.

25. Callahan, Benjamin J, Paul J McMurdie, Michael J Rosen, Andrew W Han, Amy Jo A Johnson, and Susan P Holmes. “DADA2: High-Resolution Sample Inference from Illumina Amplicon Data.” Nature Methods 13, no. 7 (July 2016): 581–83. https://doi.org/10.1038/nmeth.3869.

26. McDonald, Daniel, Morgan N Price, Julia Goodrich, Eric P Nawrocki, Todd Z DeSantis, Alexander Probst, Gary L Andersen, Rob Knight, and Philip Hugenholtz. “An Improved Greengenes Taxonomy with Explicit Ranks for Ecological and Evolutionary Analyses of Bacteria and Archaea.” The ISME Journal 6, no. 3 (March 2012): 610–18. https://doi.org/10.1038/ismej.2011.139.

27. Nilsson, Rolf Henrik, Karl-Henrik Larsson, Andy F S Taylor, Johan Bengtsson-Palme, Thomas S Jeppesen, Dmitry Schigel, Peter Kennedy, et al. “The UNITE Database for Molecular Identification of Fungi: Handling Dark Taxa and Parallel Taxonomic Classifications.” Nucleic Acids Research 47, no. D1 (January 8, 2019): D259–64. https://doi.org/10.1093/nar/gky1022.

28. Bushnell, Brian. “BBMap: A Fast, Accurate, Splice-Aware Aligner.” Lawrence Berkeley National Lab.(LBNL), 2014.

29. Cantu, Vito Adrian, Jeffrey Sadural, and Robert Edwards. “PRINSEQ++, a Multi-Threaded Tool for Fast and Efficient Quality Control and Preprocessing of Sequencing Datasets.” Preprint. PeerJ Preprints, February 27, 2019. https://doi.org/10.7287/peerj.preprints.27553v1.

30. Langmead, Ben, and Steven L Salzberg. “Fast Gapped-Read Alignment with Bowtie 2.” Nature Methods 9, no. 4 (April 2012): 357–59. https://doi.org/10.1038/nmeth.1923.

31. Nayfach, Stephen, Zhou Jason Shi, Rekha Seshadri, Katherine S. Pollard, and Nikos C. Kyrpides. “New Insights from Uncultivated Genomes of the Global Human Gut Microbiome.” Nature 568, no. 7753 (April 2019): 505–10. https://doi.org/10.1038/s41586-019-1058-x.

32. Mandal, Siddhartha, Will Van Treuren, Richard A. White, Merete Eggesbø, Rob Knight, and Shyamal D. Peddada. “Analysis of Composition of Microbiomes: A Novel Method for Studying Microbial Composition.” Microbial Ecology in Health & Disease 26, no. 0 (May 29, 2015). https://doi.org/10.3402/mehd.v26.27663.

33. Ignatiadis, Nikolaos, Bernd Klaus, Judith B Zaugg, and Wolfgang Huber. “Data-Driven Hypothesis Weighting Increases Detection Power in Genome-Scale Multiple Testing.” Nature Methods 13, no. 7 (July 2016): 577–80. https://doi.org/10.1038/nmeth.3885.

34. Beghini, Francesco, Lauren J McIver, Aitor Blanco-Míguez, Leonard Dubois, Francesco Asnicar, Sagun Maharjan, Ana Mailyan, et al. “Integrating Taxonomic, Functional, and Strain-Level Profiling of Diverse Microbial Communities with BioBakery 3.” ELife 10 (May 4, 2021): e65088. https://doi.org/10.7554/eLife.65088.

35. Li, Dinghua, Ruibang Luo, Chi-Man Liu, Chi-Ming Leung, Hing-Fung Ting, Kunihiko Sadakane, Hiroshi Yamashita, and Tak-Wah Lam. “MEGAHIT v1.0: A Fast and Scalable Metagenome Assembler Driven by Advanced Methodologies and Community Practices.” Methods 102 (June 2016): 3–11. https://doi.org/10.1016/j.ymeth.2016.02.020.

36. Hyatt, Doug, Gwo-Liang Chen, Philip F LoCascio, Miriam L Land, Frank W Larimer, and Loren J Hauser. “Prodigal: Prokaryotic Gene Recognition and Translation Initiation Site Identification.” BMC Bioinformatics 11, no. 1 (December 2010): 119. https://doi.org/10.1186/1471-2105-11-119.

37. Huerta-Cepas, Jaime, Damian Szklarczyk, Davide Heller, Ana Hernández-Plaza, Sofia K Forslund, Helen Cook, Daniel R Mende, et al. “EggNOG 5.0: A Hierarchical, Functionally and Phylogenetically Annotated Orthology Resource Based on 5090 Organisms and 2502 Viruses.” Nucleic Acids Research 47, no. D1 (January 8, 2019): D309–14. https://doi.org/10.1093/nar/gky1085.

38. Nayfach, Stephen, and Katherine S Pollard. “Average Genome Size Estimation Improves Comparative Metagenomics and Sheds Light on the Functional Ecology of the Human Microbiome.” Genome Biology 16, no. 1 (December 2015): 51. https://doi.org/10.1186/s13059-015-0611-7.

39. Wu, G. D., J. Chen, C. Hoffmann, K. Bittinger, Y.-Y. Chen, S. A. Keilbaugh, M. Bewtra, et al. “Linking Long-Term Dietary Patterns with Gut Microbial Enterotypes.” Science 334, no. 6052 (October 7, 2011): 105–8. https://doi.org/10.1126/science.1208344.

40. Turnbaugh, P. J., V. K. Ridaura, J. J. Faith, F. E. Rey, R. Knight, and J. I. Gordon. “The Effect of Diet on the Human Gut Microbiome: A Metagenomic Analysis in Humanized Gnotobiotic Mice.” Science Translational Medicine 1, no. 6 (November 11, 2009): 6ra14–6ra14. https://doi.org/10.1126/scitranslmed.3000322.

41. Hall, Andrew Brantley, Moran Yassour, Jenny Sauk, Ashley Garner, Xiaofang Jiang, Timothy Arthur, Georgia K. Lagoudas, et al. “A Novel Ruminococcus Gnavus Clade Enriched in Inflammatory Bowel Disease Patients.” Genome Medicine 9, no. 1 (November 28, 2017): 103. https://doi.org/10.1186/s13073-017-0490-5.

42. Haghi, Fakhri, Elshan Goli, Bahman Mirzaei, and Habib Zeighami. “The Association between Fecal Enterotoxigenic B. Fragilis with Colorectal Cancer.” BMC Cancer 19, no. 1 (December 2019): 879. https://doi.org/10.1186/s12885-019-6115-1.

43. Ulger Toprak, N., A. Yagci, B.M. Gulluoglu, M.L. Akin, P. Demirkalem, T. Celenk, and G. Soyletir. “A Possible Role of Bacteroides Fragilis Enterotoxin in the Aetiology of Colorectal Cancer.” Clinical Microbiology and Infection 12, no. 8 (August 2006): 782–86. https://doi.org/10.1111/j.1469-0691.2006.01494.x.

44. Cheng, Wai Teng, Haresh Kumar Kantilal, and Fabian Davamani. “The Mechanism of Bacteroides Fragilis Toxin Contributes to Colon Cancer Formation.” The Malaysian Journal of Medical Sciences: MJMS 27, no. 4 (July 2020): 9–21. https://doi.org/10.21315/mjms2020.27.4.2.

45. Ridlon, Jason M., and Phillip B. Hylemon. “Identification and Characterization of Two BileAcid Coenzyme A Transferases from Clostridium Scindens, a Bile Acid 7α-Dehydroxylating Intestinal Bacterium.” Journal of Lipid Research 53, no. 1 (January 2012): 66–76. https://doi.org/10.1194/jlr.M020313.

46. Marion, Solenne, Nicolas Studer, Lyne Desharnais, Laure Menin, Stéphane Escrig, Anders Meibom, Siegfried Hapfelmeier, and Rizlan Bernier-Latmani. “In Vitro and in Vivo Characterization of Clostridium Scindens Bile Acid Transformations.” Gut Microbes 10, no. 4 (July 4, 2019): 481–503. https://doi.org/10.1080/19490976.2018.1549420.

47. Ajouz, Hana, Deborah Mukherji, and Ali Shamseddine. “Secondary Bile Acids: An Underrecognized Cause of Colon Cancer.” World Journal of Surgical Oncology 12, no. 1 (2014): 164. https://doi.org/10.1186/1477-7819-12-164.

48. Ocvirk, Soeren, and Stephen J.D. O’Keefe. “Dietary Fat, Bile Acid Metabolism and Colorectal Cancer.” Seminars in Cancer Biology, October 2020, S1044579X2030208X. https://doi.org/10.1016/j.semcancer.2020.10.003.

49. Goncalves, Marcus D., Changyuan Lu, Jordan Tutnauer, Travis E. Hartman, Seo-Kyoung Hwang, Charles J Murphy, Chantal Pauli, et al. “High-Fructose Corn Syrup Enhances Intestinal Tumor Growth in Mice.” Science 363, no. 6433 (March 22, 2019): 1345–49. https://doi.org/10.1126/science.aat8515.

50. Michaud, Dominique S., Charles S. Fuchs, Simin Liu, Walter C. Willett, Graham A. Colditz, and Edward Giovannucci. “Dietary Glycemic Load, Carbohydrate, Sugar, and Colorectal Cancer Risk in Men and Women.” Cancer Epidemiology, Biomarkers & Prevention: A Publication of the American Association for Cancer Research, Cosponsored by the American Society of Preventive Oncology 14, no. 1 (January 2005): 138–47.

51. Lundt, Inge, and Shukun Yu. “1,5-Anhydro-d-Fructose: Biocatalytic and Chemical Synthetic Methods for the Preparation, Transformation and Derivatization.” Carbohydrate Research 345, no. 2 (January 2010): 181–90. https://doi.org/10.1016/j.carres.2009.11.004.

52. Ito, Takashi, Takaaki Totoki, Seiya Takada, Shotaro Otsuka, and Ikuro Maruyama. “Potential Roles of 1,5-Anhydro-d-Fructose in Modulating Gut Microbiome in Mice.” Scientific Reports 11, no. 1 (December 2021): 19648. https://doi.org/10.1038/s41598-021-99052-y.

53. Meng, Xiaojie, Ko-ichi Kawahara, Yuko Nawa, Naoki Miura, Binita Shrestha, Salunya Tancharoen, Hisayo Sameshima, Teruto Hashiguchi, and Ikuro Maruyama. “1,5-Anhydro-d-Fructose Attenuates Lipopolysaccharide-Induced Cytokine Release via Suppression of NF-ΚB P65 Phosphorylation.” Biochemical and Biophysical Research Communications 380, no. 2 (March 2009): 343–48. https://doi.org/10.1016/j.bbrc.2009.01.084.

54. Jin, Ye, Yang Liu, Lei Zhao, Fuya Zhao, Jing Feng, Shengda Li, Huinan Chen, et al. “Gut Microbiota in Patients after Surgical Treatment for Colorectal Cancer.” Environmental Microbiology 21, no. 2 (February 2019): 772–83. https://doi.org/10.1111/1462-2920.14498.

55. Gholizadeh, Pourya, Hosein Eslami, and Hossein Samadi Kafil. “Carcinogenesis Mechanisms of Fusobacterium Nucleatum.” Biomedicine & Pharmacotherapy 89 (May 2017): 918–25. https://doi.org/10.1016/j.biopha.2017.02.102.

56. Bess, Elizabeth N., Jordan E. Bisanz, Fauna Yarza, Annamarie Bustion, Barry E. Rich, Xingnan Li, Seiya Kitamura, et al. “Genetic Basis for the Cooperative Bioactivation of Plant Lignans by Eggerthella Lenta and Other Human Gut Bacteria.” Nature Microbiology 5, no. 1 (January 2020): 56–66. https://doi.org/10.1038/s41564-019-0596-1.

57. Webb, Amy L., and Marjorie L. McCullough. “Dietary Lignans: Potential Role in Cancer Prevention.” Nutrition and Cancer 51, no. 2 (March 2005): 117–31. https://doi.org/10.1207/s15327914nc5102_1.

58. Adlercreutz, Herman. “Lignans and Human Health.” Critical Reviews in Clinical Laboratory Sciences 44, no. 5–6 (January 2007): 483–525. https://doi.org/10.1080/10408360701612942.

59. Aune, D., D. S. M. Chan, R. Lau, R. Vieira, D. C. Greenwood, E. Kampman, and T. Norat. “Dietary Fibre, Whole Grains, and Risk of Colorectal Cancer: Systematic Review and Dose-Response Meta-Analysis of Prospective Studies.” BMJ 343, no. nov10 1 (November 11, 2011): d6617–d6617. https://doi.org/10.1136/bmj.d6617.

60. Donohoe, Dallas R., Nikhil Garge, Xinxin Zhang, Wei Sun, Thomas M. O’Connell, Maureen K. Bunger, and Scott J. Bultman. “The Microbiome and Butyrate Regulate Energy Metabolism and Autophagy in the Mammalian Colon.” Cell Metabolism 13, no. 5 (May 4, 2011): 517–26. https://doi.org/10.1016/j.cmet.2011.02.018.

61. Hague, Angela, Douglas J. E. Elder, Diane J. Hicks, and Christos Paraskeva. “Apoptosis in Colorectal Tumour Cells: Induction by the Short Chain Fatty Acids Butyrate, Propionate and Acetate and by the Bile Salt Deoxycholate.” International Journal of Cancer 60, no. 3 (January 27, 1995): 400–406. https://doi.org/10.1002/ijc.2910600322.

62. Hamer, H. M., D. Jonkers, K. Venema, S. Vanhoutvin, F. J. Troost, and R.-J. Brummer. “Review Article: The Role of Butyrate on Colonic Function.” Alimentary Pharmacology & Therapeutics 27, no. 2 (October 26, 2007): 104–19. https://doi.org/10.1111/j.1365-2036.2007.03562.x.

63. Davie, James R. “Inhibition of Histone Deacetylase Activity by Butyrate.” The Journal of Nutrition 133, no. 7 (July 1, 2003): 2485S–2493S. https://doi.org/10.1093/jn/133.7.2485S.

64. Goldstein, Neal S. “Serrated Pathway and APC (Conventional)-Type Colorectal Polyps: Molecular-Morphologic Correlations, Genetic Pathways, and Implications for Classification.” American Journal of Clinical Pathology 125, no. 1 (January 2006): 146–53. https://doi.org/10.1309/87BD0C6UCGUG236J.

