## Supplemental Figures and Tables for "Distinct Colon Mucosa Microbiomes associated with Tubular Adenomas and Serrated Polyps"

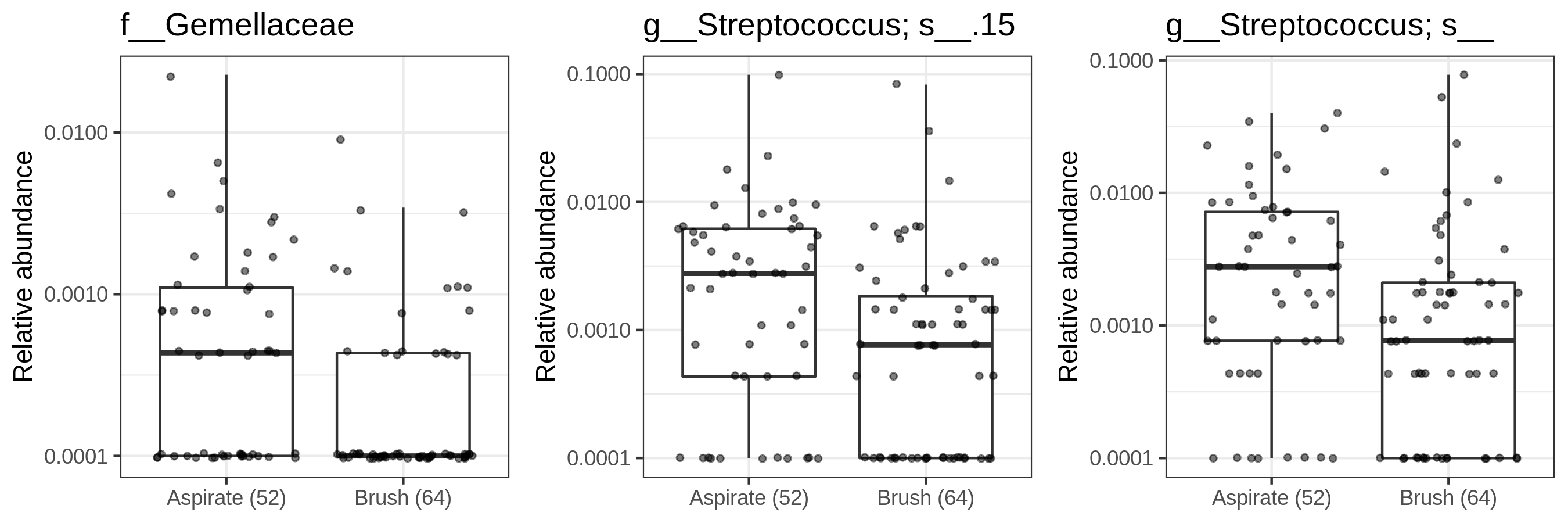

***Supplemental Figure 1:*** Box plots displaying the relative abundance of microbes determined to be differentially abundant by ANCOM2. Data is from mucosal brushes and mucosal aspirates from the first sample set. Each point is one sample, with multiple samples per individual. Plots are labeled with the most specific taxonomic rank for each ASV. A pseudo-count of 0.0001 was added to visualize samples which had relative abundances of zero, since the y axis is scaled to log_10_. The center line within each box defines the median, boxes define the upper and lower quartiles, and whiskers define 1.5x the interquartile range.

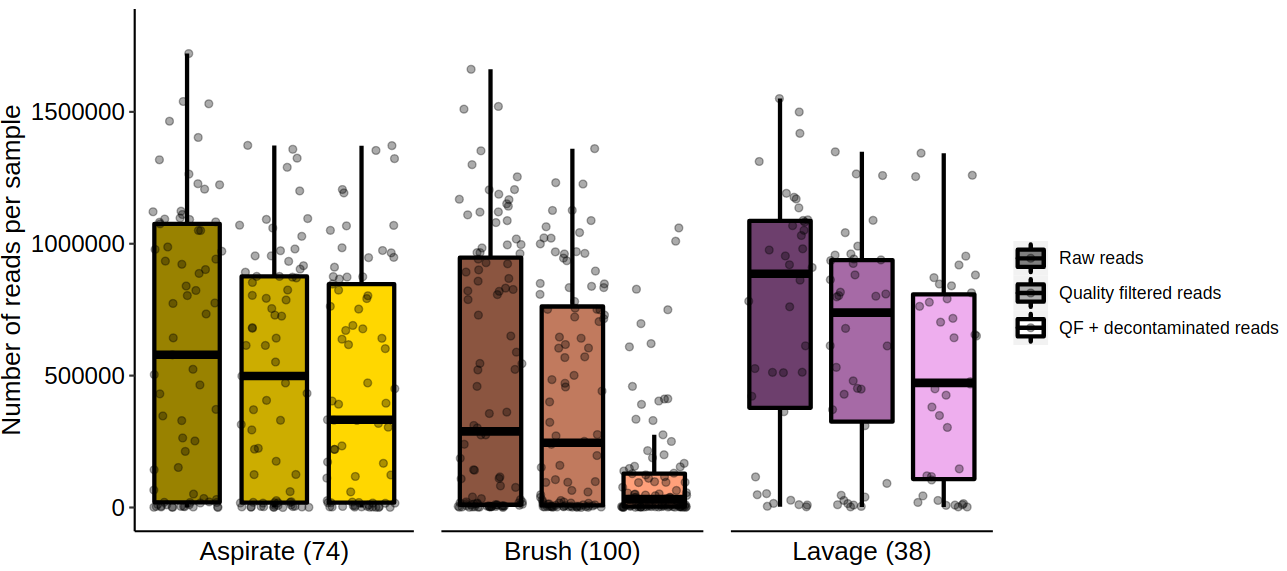

***Supplemental Figure 2:*** Box plots showing the number of reads per sample produced by a pilot shotgun sequencing run using mucosal brushes, mucosal aspirates, and lavage aliquots from the first sample set. Each point is one sample, with multiple samples per individual. The number of samples per sampling method is denoted parenthetically. The center line within each box defines the median, boxes define the upper and lower quartiles, and whiskers define 1.5x the interquartile range. ‘Raw reads’ refers to the number of reads produced by the Illumina NextSeq platform. ‘Quality filtered reads’ refers to the number of reads after removing reads with a quality score lower than a mean of 28. ‘QF + decontaminated reads’ refers to the number of reads after removing human-derived reads.

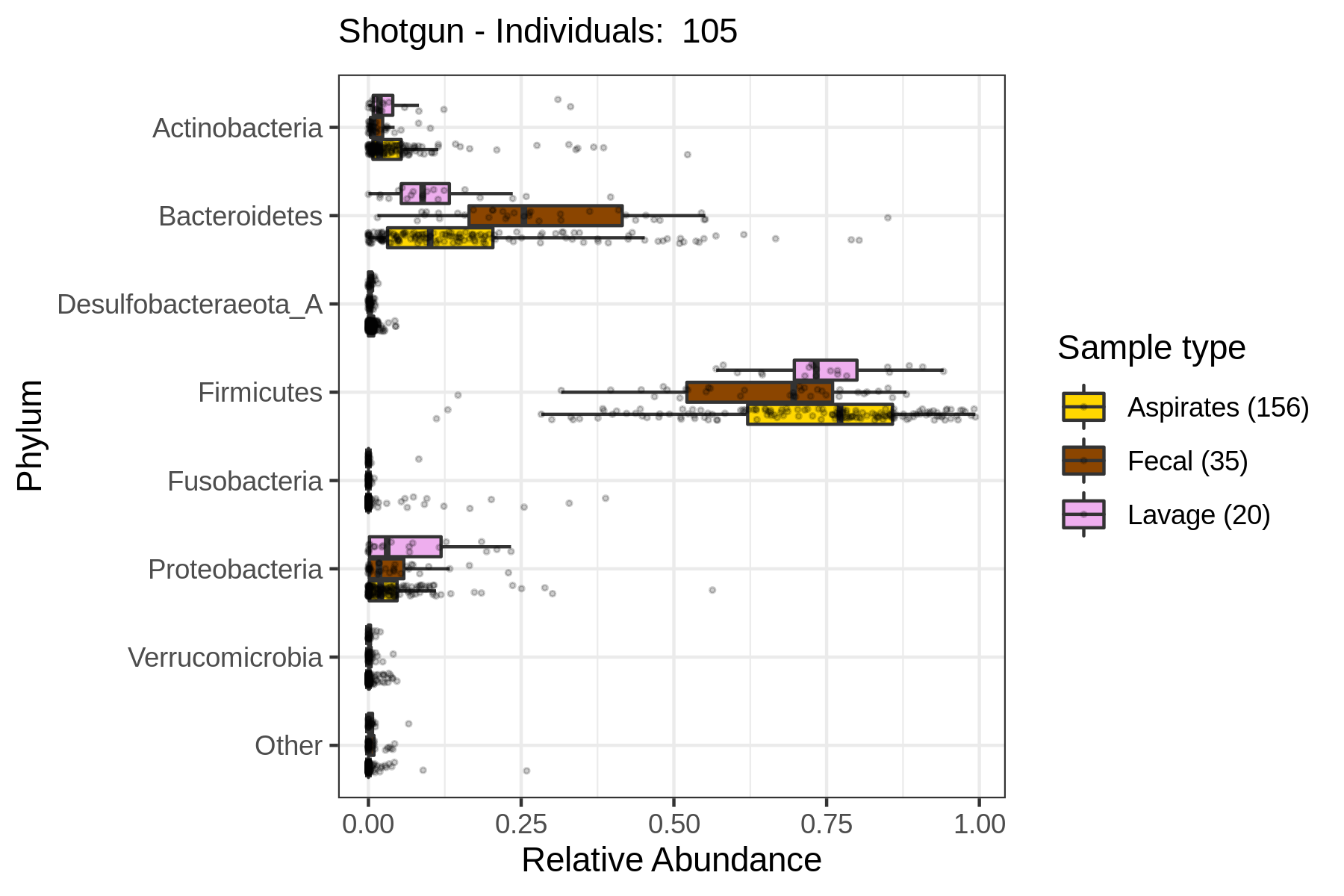

***Supplemental Figure 3:*** Box plots showing the relative abundance of the top seven most abundant microbial phyla across mucosal aspirates, lavage aliquots, and fecal samples from the second sample. Each point is one sample, with multiple samples per individual. The number of samples per sampling method is denoted parenthetically. The center line within each box defines the median, boxes define the upper and lower quartiles, and whiskers define 1.5x the interquartile range.

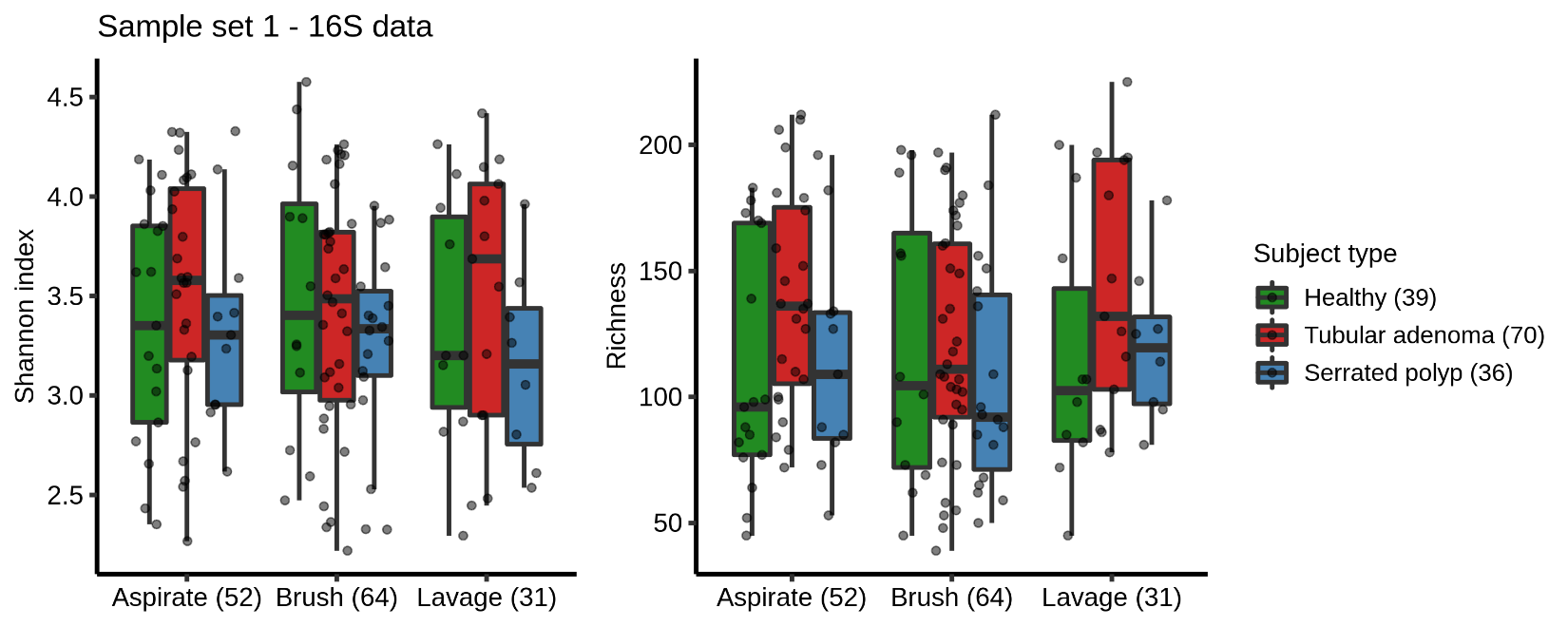

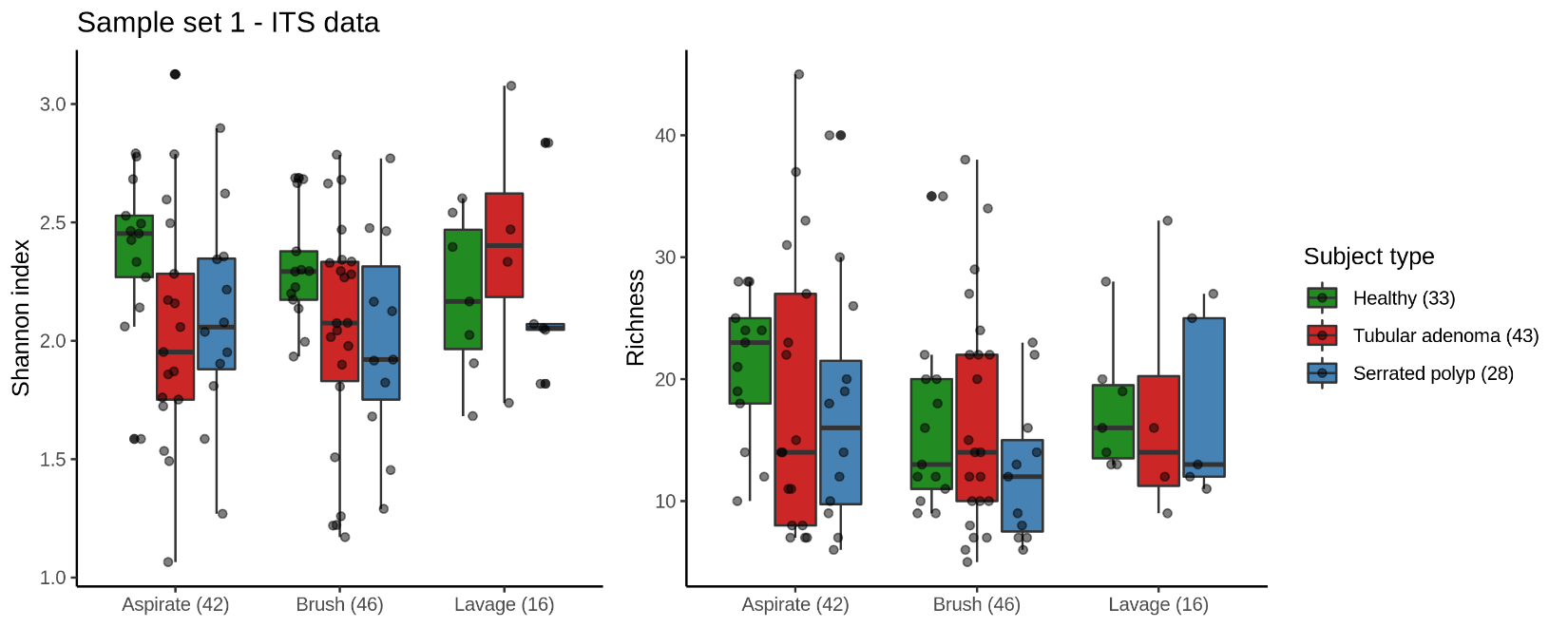

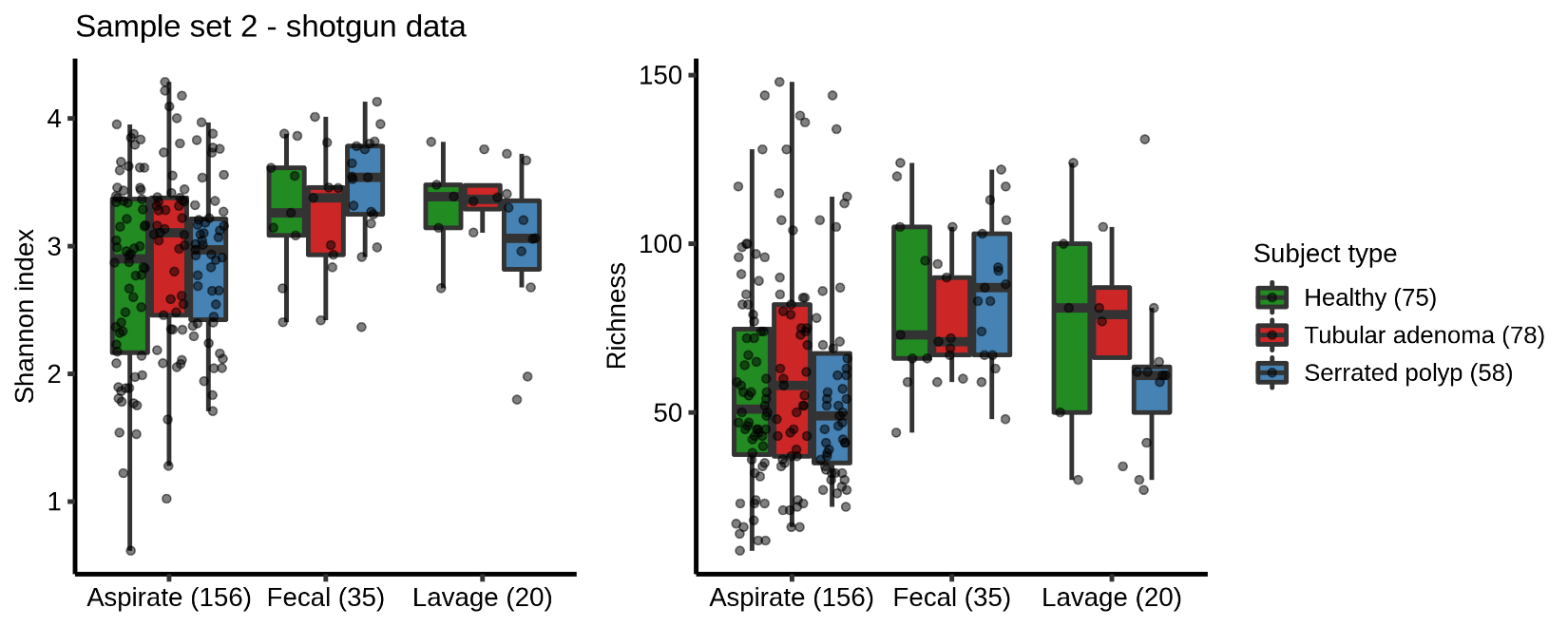

***Supplemental Figure 4:*** Box plots showing Shannon diversity and richness estimates across the first and second sample sets. The number of samples for each sampling method and subject type are denoted parenthetically. Each point is one sample, with multiple samples per individual. The center line within each box defines the median, boxes define the upper and lower quartiles, and whiskers define 1.5x the interquartile range. There was significantly increased Shannon diversity in healthy ITS samples (Linear mixed effects model: p = 0.03)

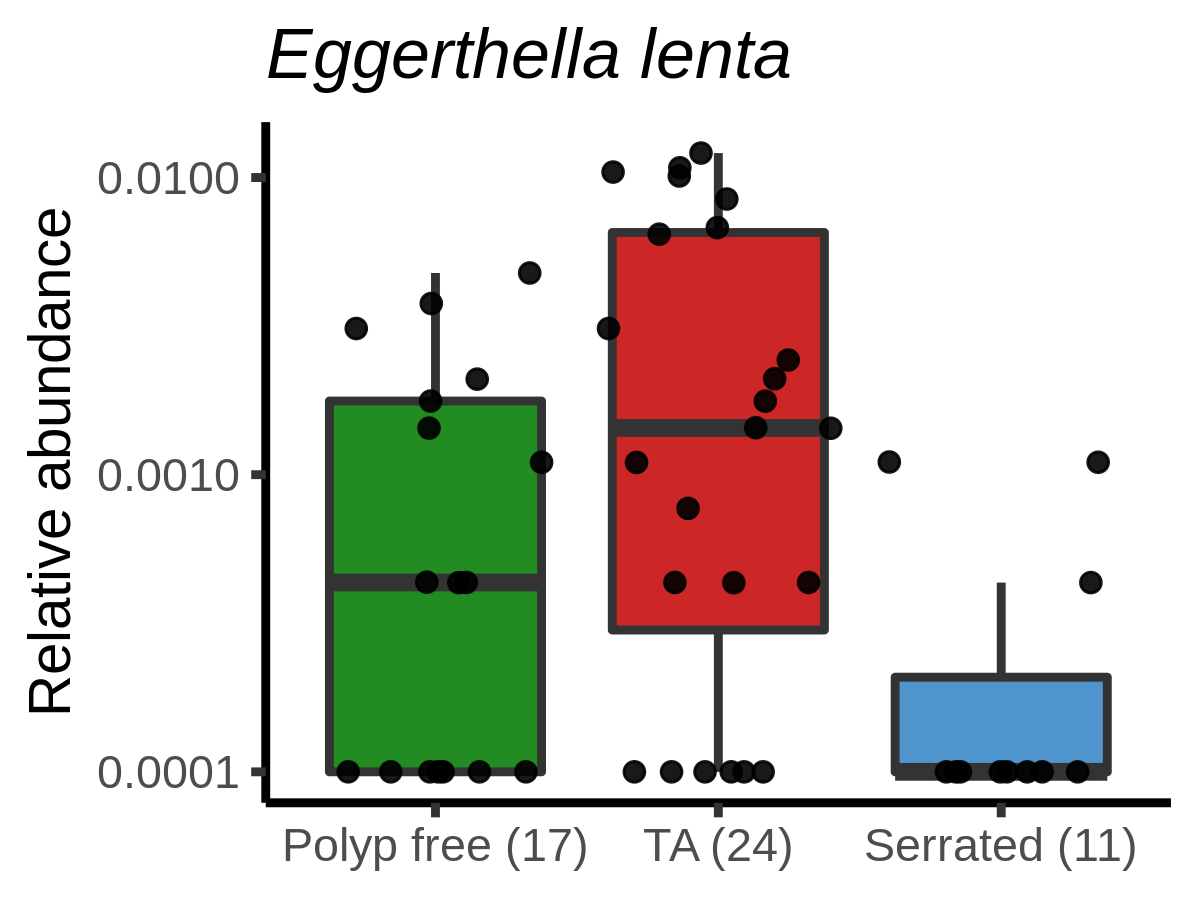

***Supplemental Figure 5:*** A box plot showing the relative abundance of *E. lenta* in 16S mucosal aspirates from the first sample set across subject types. A pseudo-count of 0.0001 was added to visualize samples which had a relative abundance of zero, since the y axis is scaled to log_10_. The center line within each box defines the median, boxes define the upper and lower quartiles, and whiskers define 1.5x the interquartile range.

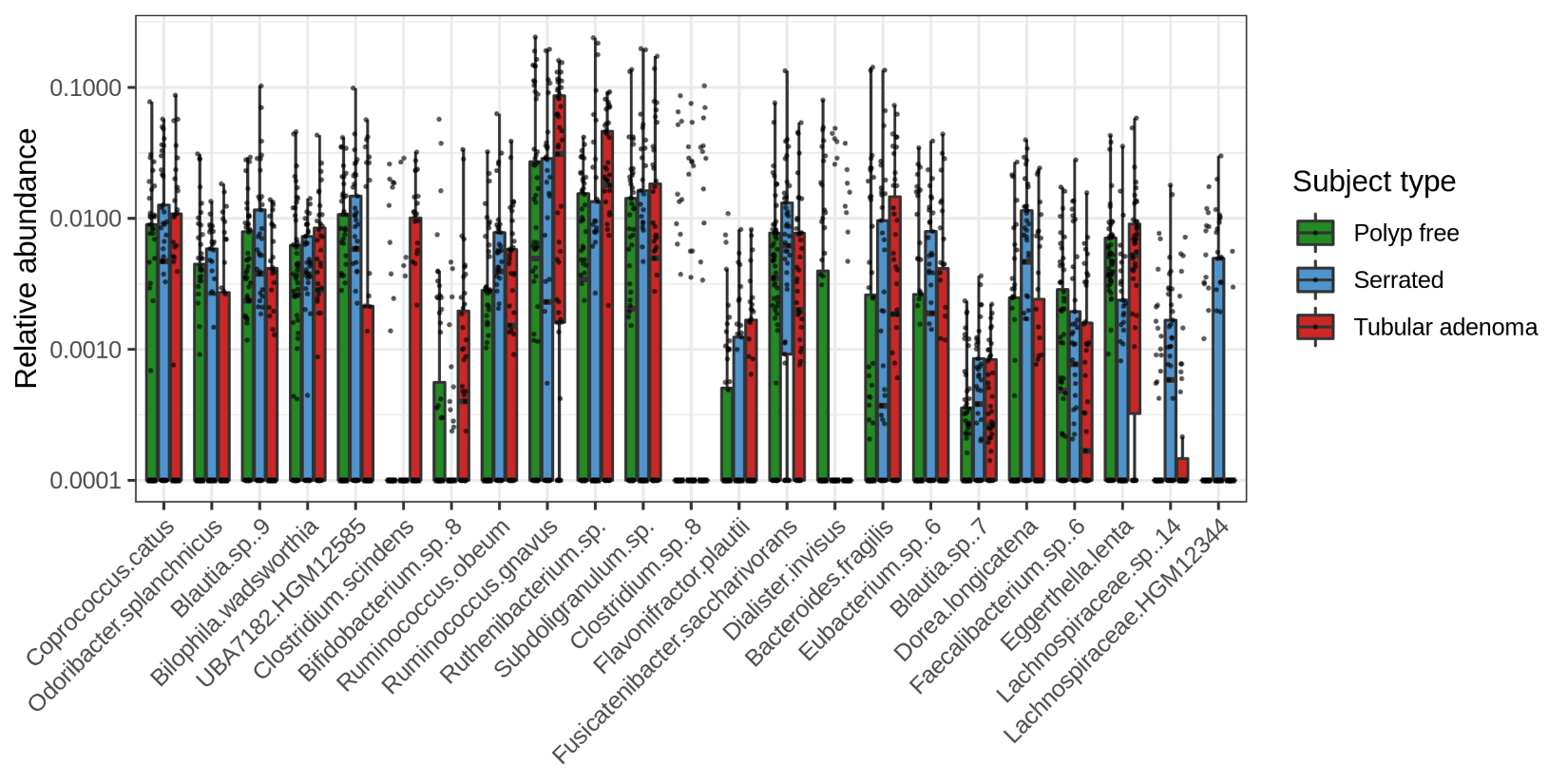

***Supplemental Figure 6:*** Box plots displaying the relative abundances of the top variables of importance as determined by Random Forest. Only mucosal aspirates from the second sample set are included. Each point is one sample, with multiple samples per individual. A pseudo-count of 0.0001 was added to visualize samples which had relative abundances of zero, since the y axis is scaled to log_10_. The center line within each box defines the median, boxes define the upper and lower quartiles, and whiskers define 1.5x the interquartile range.

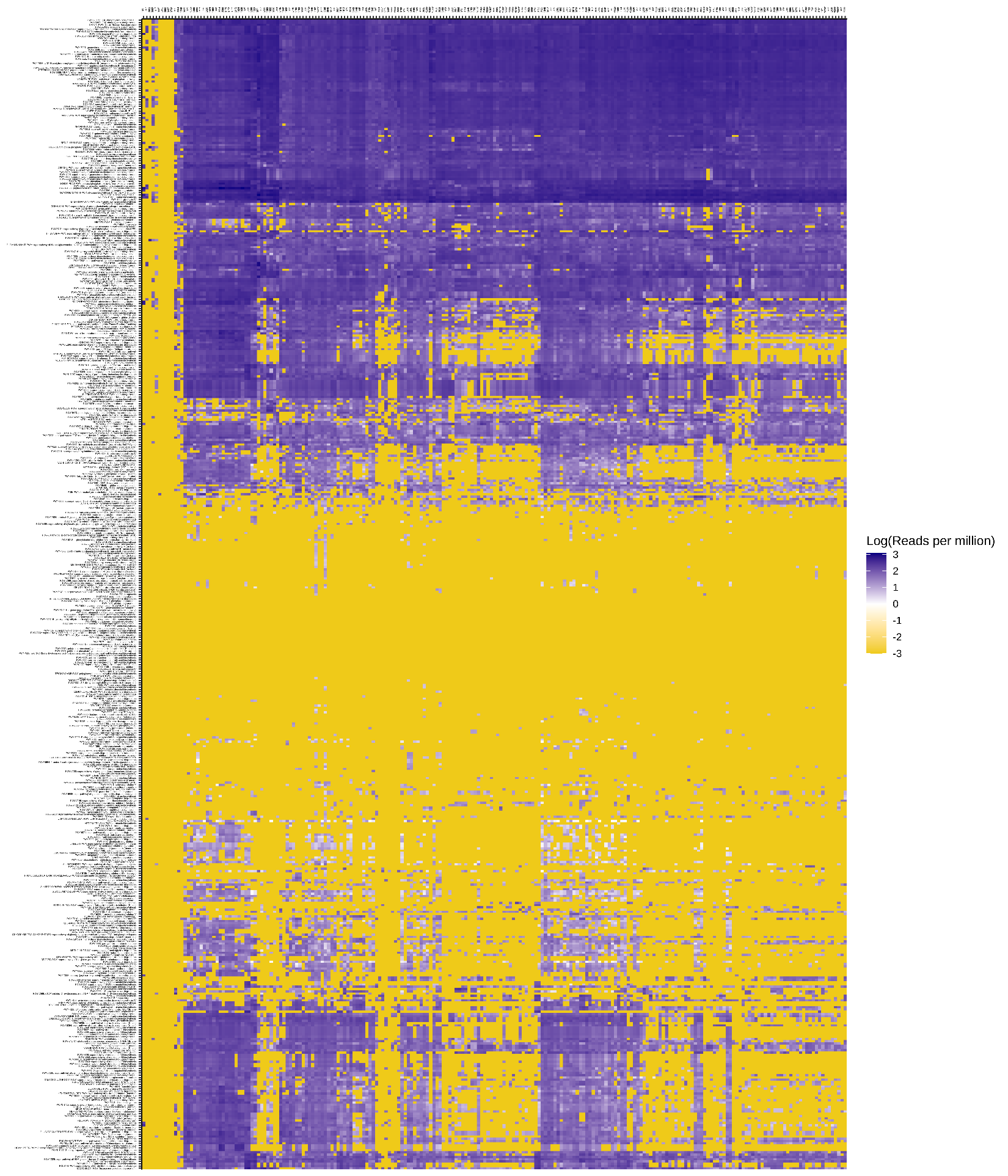

***Supplemental Figure 7:*** A heatmap displaying the abundance of microbial pathways from the second sample set. Abundances are visualized in log_10_(Reads per million). A total 507 pathways were identified.

|  | Polyp-free | TA-bearing | SP-bearing (HPP/SSP) | Unknown or Other | Total |
| --- | --- | --- | --- | --- | --- |
| Mucosal brushes [On-polyp] | 197 [0] | 168 [58] | 112 (61 [18]/ 51 [17]) | 48 [14] | 525 |
| Mucosal aspirates | 350 | 280 | 195 (111/84) | 72 | 897 |
| Lavage aspirates | 159 | 135 | 93 (54/39) | 36 | 423 |
| Fecal samples | 9 | 17 | 9 (6/3) | 3 | 38 |
| Total | 715 | 600 | 409 | 159 | 1883 |

***Supplemental Table 1:*** A table showing the number of samples collected. Across rows, the number of each sample type is listed. For mucosal brushes, the number within the bracket corresponds to the number of brush samples taken directly from polyp tissue (as opposed to brushing non-polyp tissue). Across columns, the subject type classification is given. The number of samples per hyperplastic polyps (HPP) and sessile serrated polyps (SSP) are denoted parenthetically for the SP-bearing category. Samples were collected from a total of 140 unique individuals.

| 16S | Polyp-free | TA-bearing | SP-bearing (HPP/SSP) | Unknown or Other | Total |
| --- | --- | --- | --- | --- | --- |
| Mucosal brushes [On-polyp] | 13 [0] | 31 [11] | 18 (10 [2]/8 [4]) | 2 [0] | 64 |
| Mucosal aspirates | 17 | 23 | 12 (7/5) | 0 | 52 |
| Lavage aspirates | 11 | 12 | 8 (4/4) | 0 | 31 |
| Fecal samples | 0 | 0 | 0 | 0 | 0 |
| Total | 41 | 66 | 38 | 2 | 147 |

***Supplemental Table 2:*** A table showing the number of samples with high quality sequencing reads in sample set 1, using 16S amplicon sequencing. Across rows, the number of each sample type is listed. For mucosal brushes, the number within the bracket corresponds to the number of brush samples taken directly from polyp tissue (as opposed to brushing non-polyp tissue). Across columns, the subject type classification is given. The number of samples per hyperplastic polyps (HPP) and sessile serrated polyps (SSP) are denoted parenthetically for the SP-bearing category. A total of 38 unique individuals were represented in this data.

| ITS | Polyp-free | TA-bearing | SP-bearing (HPP/SSP) | Unknown or Other | Total |
| --- | --- | --- | --- | --- | --- |
| Mucosal brushes [On-polyp] | 13 [0] | 18 [6] | 13 (6 [1]/7 [1]) | 2 | 46 |
| Mucosal aspirates | 13 | 15 | 12 (5/7) | 2 | 42 |
| Lavage aspirates | 7 | 3 | 6 (2/4) | 0 | 16 |
| Fecal samples | 0 | 0 | 0 | 0 | 0 |
| Total | 33 | 36 | 31 | 4 | 104 |

***Supplemental Table 3:*** A table showing the number of samples with high quality sequencing reads in sample set 1, using ITS amplicon sequencing. Across rows, the number of each sample type is listed. For mucosal brushes, the number within the bracket corresponds to the number of brush samples taken directly from polyp tissue (as opposed to brushing non-polyp tissue). Across columns, the subject type classification is given. The number of samples per hyperplastic polyps (HPP) and sessile serrated polyps (SSP) are denoted parenthetically for the SP-bearing category. A total of 35 unique individuals were represented in this data.

| WGS | Polyp-free | TA-bearing | SP-bearing (HPP/SSP) | Unknown or Other | Total |
| --- | --- | --- | --- | --- | --- |
| Mucosal brushes [On-polyp] | 0 | 0 | 0 | 0 | 0 |
| Mucosal aspirates | 64 | 47 | 45 (24/17) | 23 | 179 |
| Lavage aspirates | 5 | 11 | 4 (2/2) | 1 | 21 |
| Fecal samples | 9 | 17 | 9 (6/3) | 3 | 38 |
| Total | 78 | 75 | 58 | 27 | 238 |

***Supplemental Table 4:*** A table showing the number of samples with high quality sequencing reads in sample set 2, using whole-genome shotgun sequencing. The number of each sample type is listed across rows. Across columns, the subject type classification is given. Additionally, the number of samples per hyperplastic polyps (HPP) and sessile serrated polyps (SSP) are denoted parenthetically for the SP-bearing category. A total of 117 unique individuals were represented in this data.

| FACTOR | DoF | SoS | MS | F MODEL | R^2^ | P-VAL |
| --- | --- | --- | --- | --- | --- | --- |
| BMI | 1 | 1.53 | 1.53 | 15.40 | 0.03 | 0.001 |
| AGE | 1 | 1.02 | 1.02 | 10.29 | 0.02 | 0.001 |
| ETHNICITY | 4 | 5.35 | 1.34 | 13.51 | 0.11 | 0.001 |
| SEX | 1 | 1.23 | 1.23 | 12.38 | 0.02 | 0.001 |
| SUBJECT TYPE | 2 | 2.28 | 1.14 | 11.50 | 0.05 | 0.001 |
| SUBJECT TYPE: INDIVIDUAL | 28 | 28.21 | 1.00 | 10.17 | 0.56 | 0.001 |
| SUBJECT TYPE: INDIVIDUAL: SAMPLE TYPE | 63 | 6.29 | 0.10 | 1.01 | 0.12 | 0.491 |
| RESIDUALS | 46 | 4.56 | 0.10 |  | 0.09 |  |
| TOTAL | 146 | 50.46 |  |  | 1.00 |  |

PERMANOVA formula: 16S_ASV_table ~ BMI + Age + Ethnicity + Sex + Subject Type / Individual / Sample Type, strata = Plate

***Supplemental Table 5:*** PERMANOVA analysis of brushes, mucosal aspirates, and lavage aliquots from the first sample set using 16S sequencing. The distance matrix method used was Bray-Curtis dissimilarity. Subject type includes polyp-free, tubular adenoma-bearing, and serrated polyp-bearing samples. Individuals are nested within subject type, and sample type is nested within the individual.

| FACTOR | DoF | SoS | MS | F MODEL | R^2^ | P-VAL |
| --- | --- | --- | --- | --- | --- | --- |
| BMI | 1 | 0.55 | 0.55 | 1.35 | 0.01 | 0.029 |
| AGE | 1 | 0.54 | 0.54 | 1.32 | 0.01 | 0.041 |
| SEX | 1 | 0.43 | 0.43 | 1.06 | 0.01 | 0.320 |
| ETHNICITY | 3 | 1.43 | 0.48 | 1.17 | 0.03 | 0.058 |
| SUBJECT TYPE | 2 | 0.94 | 0.47 | 1.16 | 0.02 | 0.106 |
| SUBJECT TYPE: INDIVIDUAL | 25 | 12.26 | 0.49 | 1.21 | 0.28 | 0.001 |
| SUBJECT TYPE: INDIVIDUAL: SAMPLE TYPE | 40 | 17.02 | 0.43 | 1.05 | 0.38 | 0.123 |
| RESIDUALS | 28 | 11.37 | 0.41 |  | 0.26 |  |
| TOTAL | 101 | 44.53 |  |  | 1.00 |  |

PERMANOVA formula: ITS_ASV_table ~ BMI + Age + Sex + Ethnicity + Subject Type / Individual / Sample Type

***Supplemental Table 6:*** PERMANOVA analysis of brushes, mucosal aspirates, and lavage aliquots from the first sample set using ITS sequencing. The distance matrix method used was Bray-Curtis dissimilarity. Subject type includes polyp-free, tubular adenoma-bearing, and serrated polyp-bearing samples. Individuals are nested within subject type, and sample type is nested within the individual.

| FACTOR | DoF | SoS | MS | F MODEL | R^2^ | P-VAL |
| --- | --- | --- | --- | --- | --- | --- |
| BMI | 1 | 0.61 | 0.61 | 21.03 | 0.01 | 0.001 |
| AGE | 1 | 0.90 | 0.90 | 31.11 | 0.01 | 0.001 |
| ETHNICITY | 5 | 3.50 | 0.70 | 24.14 | 0.04 | 0.001 |
| SEX | 1 | 0.71 | 0.71 | 24.63 | 0.01 | 0.001 |
| SUBJECT TYPE | 2 | 1.49 | 0.77 | 25.76 | 0.02 | 0.001 |
| SUBJECT TYPE: INDIVIDUAL | 94 | 60.21 | 0.64 | 22.11 | 0.75 | 0.001 |
| SUBJECT TYPE: INDIVIDUAL: SAMPLE TYPE | 43 | 11.30 | 0.26 | 9.07 | 0.14 | 0.001 |
| RESIDUALS | 63 | 1.83 | 0.03 |  | 0.02 |  |
| TOTAL | 210 | 80.55 |  |  | 1.00 |  |

PERMANOVA formula: OTU_table ~ BMI + Age + Sex + Ethnicity + Subject Type / Individual / Sample Type, strata = Plate

***Supplemental Table 7:*** PERMANOVA analysis of mucosal aspirates, lavage aliquots, and fecal samples from the second sample set using shotgun sequencing. The distance matrix method used was Bray-Curtis dissimilarity. Subject type includes polyp-free, tubular adenoma-bearing, and serrated polyp-bearing samples. Individuals are nested within subject type, and sample type is nested within the individual.

| taxa_id | W | detected_0.9 | detected_0.8 | detected_0.7 | detected_0.6 |
| --- | --- | --- | --- | --- | --- |
| UBA1381.sp. | 134 | TRUE | TRUE | TRUE | TRUE |
| Dorea.formicigenerans | 132 | TRUE | TRUE | TRUE | TRUE |
| Ruminococcus.torques | 132 | TRUE | TRUE | TRUE | TRUE |
| Bacteroides.fragilis | 131 | TRUE | TRUE | TRUE | TRUE |
| Clostridium.ramosum | 131 | TRUE | TRUE | TRUE | TRUE |
| Ruminococcus.bicirculans | 131 | TRUE | TRUE | TRUE | TRUE |
| Ruminococcus.gnavus | 130 | TRUE | TRUE | TRUE | TRUE |
| Coprococcus.catus | 127 | TRUE | TRUE | TRUE | TRUE |
| Ruminococcus.lactaris | 126 | TRUE | TRUE | TRUE | TRUE |
| DTU089.HGM12760 | 126 | TRUE | TRUE | TRUE | TRUE |
| Clostridium.leptum | 124 | TRUE | TRUE | TRUE | TRUE |
| Oscillibacter.sp. | 124 | TRUE | TRUE | TRUE | TRUE |
| Lachnospiraceae.sp..12 | 123 | FALSE | TRUE | TRUE | TRUE |
| D16.sp..2 | 123 | FALSE | TRUE | TRUE | TRUE |
| D16.sp. | 121 | FALSE | TRUE | TRUE | TRUE |
| Eggerthella.lenta | 119 | FALSE | TRUE | TRUE | TRUE |
| Dorea.sp..2 | 119 | FALSE | TRUE | TRUE | TRUE |
| Lachnospira.sp..2 | 119 | FALSE | TRUE | TRUE | TRUE |
| Dorea.longicatena.1 | 117 | FALSE | TRUE | TRUE | TRUE |
| Tyzzerella.sp..1 | 117 | FALSE | TRUE | TRUE | TRUE |
| Eubacterium.sp..15 | 116 | FALSE | TRUE | TRUE | TRUE |
| Eubacterium.sp..9 | 115 | FALSE | TRUE | TRUE | TRUE |
| UBA7182.HGM12585 | 115 | FALSE | TRUE | TRUE | TRUE |
| Alistipes.sp..3 | 114 | FALSE | TRUE | TRUE | TRUE |
| Lachnospira.pectinoschiza | 114 | FALSE | TRUE | TRUE | TRUE |
| Eubacterium.HGM12316 | 113 | FALSE | TRUE | TRUE | TRUE |
| Roseburia.intestinalis | 112 | FALSE | TRUE | TRUE | TRUE |
| Bilophila.wadsworthia | 109 | FALSE | FALSE | TRUE | TRUE |
| Clostridium.bartlettii | 109 | FALSE | FALSE | TRUE | TRUE |
| Alistipes.putredinis | 108 | FALSE | FALSE | TRUE | TRUE |
| Flavonifractor.plautii | 108 | FALSE | FALSE | TRUE | TRUE |
| Faecalibacterium.HGM13278 | 107 | FALSE | FALSE | TRUE | TRUE |
| Lachnospiraceae.HGM11862 | 105 | FALSE | FALSE | TRUE | TRUE |
| Coprococcus.comes | 105 | FALSE | FALSE | TRUE | TRUE |
| Eubacterium.sp..6 | 104 | FALSE | FALSE | TRUE | TRUE |
| ER4.sp. | 104 | FALSE | FALSE | TRUE | TRUE |
| Faecalibacterium.sp..6 | 103 | FALSE | FALSE | TRUE | TRUE |
| Anaerostipes.hadrus | 102 | FALSE | FALSE | TRUE | TRUE |
| Parabacteroides.distasonis | 101 | FALSE | FALSE | TRUE | TRUE |
| DTU089.HGM12731 | 100 | FALSE | FALSE | TRUE | TRUE |
| Intestinimonas.butyriciproducens | 100 | FALSE | FALSE | TRUE | TRUE |
| Coprococcus.HGM12238 | 99 | FALSE | FALSE | TRUE | TRUE |
| Faecalibacterium.HGM13282 | 98 | FALSE | FALSE | TRUE | TRUE |
| Escherichia.coli | 98 | FALSE | FALSE | TRUE | TRUE |

***Supplemental Table 8:*** Table of differentially abundant OTUs across fecal samples and mucosal aspirates from the second sample set using shotgun sequencing. Significance testing was performed using ANCOM2 (FDR < 0.05), adjusting for repeated measurements. “Detected 0.7” means that the microbe was differentially abundant in 70% of comparisons, which is the minimum for a microbe to be considered differentially abundant between categories.

| taxa_id | W | detected_0.9 | detected_0.8 | detected_0.7 | detected_0.6 |
| --- | --- | --- | --- | --- | --- |
| Ruminococcus.torques | 85 | TRUE | TRUE | TRUE | TRUE |
| Dorea.formicigenerans | 83 | TRUE | TRUE | TRUE | TRUE |
| Ruminococcus.bicirculans | 80 | FALSE | TRUE | TRUE | TRUE |
| Oscillibacter.sp. | 78 | FALSE | TRUE | TRUE | TRUE |
| Coprococcus.catus | 72 | FALSE | FALSE | TRUE | TRUE |
| Lachnospiraceae.sp..12 | 68 | FALSE | FALSE | TRUE | TRUE |

***Supplemental Table 9:*** Table of differentially abundant OTUs across fecal samples and lavage aliquots from the second sample set using shotgun sequencing. Significance testing was performed using ANCOM2 (FDR < 0.05), adjusting for repeated measurements. “Detected 0.7” means that the microbe was differentially abundant in 70% of comparisons, which is the minimum for a microbe to be considered differentially abundant between categories.

| FACTOR | DoF | SoS | MS | F MODEL | R^2^ | P-VAL |
| --- | --- | --- | --- | --- | --- | --- |
| COLON LOCATION | 1 | 0.69 | 0.69 | 5.08 | 0.16 | 0.005 |
| SUBJECT TYPE | 1 | 0.42 | 0.42 | 3.07 | 0.10 | 0.030 |
| SUBJECT TYPE: INDIVIDUAL | 4 | 2.28 | 0.57 | 4.17 | 0.52 | 0.004 |
| SUBJECT TYPE: INDIVIDUAL: TISSUE SITE | 6 | 0.82 | 0.14 | 0.99 | 0.19 | 0.528 |
| RESIDUALS | 1 | 0.14 | 0.14 |  | 0.03 |  |
| TOTAL | 13 | 4.35 |  |  | 1.00 |  |

PERMANOVA formula: 16S_brushes_ASV_table ~ Colon location + Subject type / Individual / Tissue site

***Supplemental Table 10:*** PERMANOVA analysis of brushes from the first sample set using 16S sequencing. The distance matrix method used was Bray-Curtis dissimilarity. Subject type includes tubular adenoma-bearing and serrated polyp-bearing samples. Tissue site includes polyp and healthy, opposite wall brushes. Individuals are nested within subject type, and tissue site is nested within the individual.

| FACTOR | DoF | SoS | MS | F MODEL | R^2^ | P-VAL |
| --- | --- | --- | --- | --- | --- | --- |
| SUBJECT TYPE | 2 | 0.75 | 0.38 | 1.02 | 0.11 | 0.467 |
| PREP TYPE | 2 | 0.80 | 0.40 | 1.08 | 0.11 | 0.332 |
| AGE | 1 | 0.34 | 0.34 | 0.92 | 0.05 | 0.670 |
| BMI | 1 | 0.36 | 0.36 | 0.97 | 0.05 | 0.381 |
| SEX | 2 | 0.67 | 0.34 | 0.91 | 0.10 | 0.760 |
| EHTNICITY | 2 | 0.70 | 0.35 | 0.95 | 0.10 | 0.678 |
| RESIDUALS | 9 | 3.32 | 0.37 |  | 0.48 |  |
| TOTAL | 19 | 6.95 |  |  | 1.00 |  |

PERMANOVA formula: Lavage_OTU_table ~ Subject type + Prep type + Age + BMI + Sex + Ethnicity, strata = Plate

***Supplemental Table 11:*** PERMANOVA analysis of lavage aliquots from the second sample set using shotgun sequencing. The distance matrix method used was Bray-Curtis dissimilarity. Subject type includes polyp-free, tubular adenoma-bearing and serrated polyp-bearing samples. Prep type refers to the laxative used during colonoscopy prep.

| FACTOR | DoF | SoS | MS | F MODEL | R^2^ | P-VAL |
| --- | --- | --- | --- | --- | --- | --- |
| SUBJECT TYPE | 2 | 0.85 | 0.43 | 1.16 | 0.07 | 0.103 |
| AGE | 1 | 0.30 | 0.30 | 0.82 | 0.02 | 0.879 |
| BMI | 1 | 0.48 | 0.48 | 1.30 | 0.04 | 0.048 |
| SEX | 2 | 0.71 | 0.35 | 0.96 | 0.06 | 0.575 |
| ETHNICITY | 3 | 1.02 | 0.34 | 0.93 | 0.08 | 0.778 |
| RESIDUALS | 25 | 9.12 | 0.38 |  | 0.73 |  |
| TOTAL | 34 | 12.55 |  |  | 1.00 |  |

PERMANOVA formula: Fecal_OTU_table ~ Subject type + Age + BMI + Sex + Ethnicity, strata = Plate

***Supplemental Table 12:*** PERMANOVA analysis of fecal samples from the second sample set using shotgun sequencing. The distance matrix method used was Bray-Curtis dissimilarity. Subject type includes polyp-free, tubular adenoma-bearing and serrated polyp-bearing samples. Prep type refers to the solution used during colonoscopy prep.

| FACTOR | DoF | SoS | MS | F MODEL | R^2^ | P-VAL |
| --- | --- | --- | --- | --- | --- | --- |
| BMI | 1 | 0.008 | 0.008 | 36.546 | 0.024 | 0.001 |
| AGE | 1 | 0.003 | 0.003 | 12.465 | 0.008 | 0.001 |
| ETHNICITY | 5 | 0.015 | 0.003 | 14.357 | 0.047 | 0.001 |
| SEX | 1 | 0.001 | 0.001 | 4.392 | 0.003 | 0.005 |
| SUBJECT TYPE | 2 | 0.005 | 0.002 | 11.993 | 0.016 | 0.001 |
| SUBJECT TYPE: INDIVIDUAL | 94 | 0.237 | 0.003 | 12.234 | 0.751 | 0.001 |
| SUBJECT TYPE: INDIVIDUAL: SAMPLE TYPE | 44 | 0.035 | 0.001 | 3.819 | 0.109 | 0.001 |
| RESIDUALS | 64 | 0.013 | 0.000 |  | 0.042 |  |
| TOTAL | 212 | 0.316 |  |  | 1.000 |  |

PERMANOVA formula: Gene_table ~ BMI + Age + Ethnicity + Sex + Subject type / Individual / Sample type, strata = Plate

***Supplemental Table 13:*** PERMANOVA analysis of functional genes within mucosal aspirates, lavage aliquots, and fecal samples from the second sample set using shotgun sequencing. The distance matrix method used was Bray-Curtis dissimilarity. Subject type includes polyp-free, tubular adenoma-bearing, and serrated polyp-bearing samples. Individuals are nested within subject type, and sample type is nested within the individual.
